# Dysregulation of the p53 pathway provides a therapeutic target in aggressive pediatric sarcomas with stem-like traits

**DOI:** 10.1101/2024.01.17.576012

**Authors:** Lucie Curylova, Iva Staniczkova Zambo, Jakub Neradil, Michal Kyr, Nicola Jurackova, Sarka Pavlova, Kristyna Polaskova, Peter Mudry, Jaroslav Sterba, Renata Veselska, Jan Skoda

**Author notes:** **Corresponding Author:** Dr. Jan Skoda, Department of Experimental Biology, Faculty of Science, Masaryk University, University Campus Bohunice - C13.326, Kamenice 5, 625 00 Brno, Czech Republic. **Competing interests:** The authors declare no competing financial interests.

## Abstract

Pediatric sarcomas are bone and soft tissue tumors that often exhibit high metastatic potential and refractory stem-like phenotypes, resulting in poor outcomes. Aggressive sarcomas frequently harbor a disrupted p53 pathway. However, whether sarcoma stemness is associated with abrogated p53 function and might be attenuated via p53 reactivation remains unclear. Here, we show that highly tumorigenic stem-like sarcoma cells exhibit dysregulated p53, making them vulnerable to drugs that restore wild-type p53 activity. Immunohistochemistry of mouse xenografts and human tumor tissues revealed that p53 dysregulations together with enhanced expression of the stemness-related transcription factors SOX2 or KLF4 are crucial features in pediatric osteosarcoma, rhabdomyosarcoma, and Ewing’s sarcoma development. p53 dysregulation appears to be an important step for sarcoma cells to acquire a fully stem-like phenotype, and p53-positive pediatric sarcomas exhibit a high frequency of early metastasis. Importantly, p53 signaling reactivation via MDM2/MDMX inhibition selectively induces apoptosis in aggressive stem-like Ewing’s sarcoma cells while sparing healthy fibroblasts. Collectively, our results suggest that restoration of canonical p53 activity provides a promising strategy for improving the treatment of pediatric sarcomas with unfavorable stem-like traits.

**HIGHLIGHTS:**

- SOX2 and KLF4 are crucial factors in pediatric sarcoma tumorigenesis
- Dysregulated p53 pathway predisposes sarcoma cells to acquire stem-like features
- p53 positivity is associated with early metastasis in pediatric sarcoma patients
- Restoring wild-type p53 signaling selectively kills stem-like Ewing’s sarcoma cells

## 1. INTRODUCTION

Sarcomas are common pediatric tumors, accounting for ∼15% of all malignancies diagnosed in children. These soft tissue and bone tumors most likely originate from dysfunctional mesenchymal stem cells and their derivatives [1,2]. Despite intensive multimodal therapy, approximately one-third of pediatric sarcoma patients experience disease recurrence, therapy resistance and metastasis—traits commonly linked to cancer stemness—which results in a dismal survival rate of only 20% [3,4]. Elucidating the mechanisms underlying the development of these aggressive therapy-resistant tumors is therefore crucial for improving the therapeutic management of pediatric sarcomas.

In addition to the expression of several putative markers [3,5–7], the expression of key pluripotency-related transcription factors (TFs), i.e., SOX2, OCT4, NANOG, KLF4 and c-MYC [8], has been linked to the stem-like phenotype of sarcoma cells [9–13]. Upregulation of these TFs is well known to alter cell proliferation, differentiation and plasticity [8,14], highlighting their roles as core functional regulators of the stem-like state and potential mediators of sarcoma tumorigenesis. Nevertheless, whether the expression levels of these TFs, either individually or in combination, might serve as predictors of pediatric sarcoma patient outcomes remains unclear.

p53 is primarily recognized as a crucial tumor suppressor and was recently identified as a regulator of stemness maintenance and differentiation in healthy and tumor tissues [1,15]. This factor also restricts cancer stem cell (CSC) self-renewal and inhibits tumor initiation and progression [15]. Because *TP53* mutations and other p53 pathway alterations (*e.g.*, *MDM2* amplification) are observed in most human sarcomas [1], targeting p53 might be a strategy for suppressing sarcoma stemness. Several approaches for restoring wild-type p53 (wt-p53) function have been designed, including reactivation or depletion of mutant p53 (mut-p53) and inhibition of p53 negative regulators from the MDM family [1]. However, whether p53 dysregulation is related to the stemness of pediatric sarcomas is unknown.

This study aimed to elucidate the potential connection between the dysregulated stemness-related transcriptional network and p53 alterations in pediatric sarcomas. Furthermore, we investigated whether restoration of normal p53 activity might effectively eliminate sarcoma CSCs. Using a unique panel of established and patient-derived osteosarcoma (OS), rhabdomyosarcoma (RMS) and Ewing’s sarcoma (ES) cell lines, we report strong evidence showing that p53 dysregulation is linked to upregulated expression of selected stemness-related TFs and aggressive tumorigenic features of pediatric sarcoma cells. Interestingly, p53 pathway dysregulation was identified as the most consistent factor contributing to development of the stem-like, tumorigenic phenotype of sarcoma cells. Consistently, immunohistochemistry analysis of samples from a cohort of pediatric OS, RMS and ES patients confirmed that p53 positivity is associated with metastasis risk. Corroborating these findings, this work provides the first insights into the therapeutic potential of attenuating sarcoma stemness by restoring p53 activity. The restoration of p53 activity preferentially affected stem-like ES cells compared with their nontumorigenic counterparts or healthy fibroblasts. Together, our findings indicate that reactivation of the dysregulated p53 pathway represents a clinically relevant strategy for improving the treatment of aggressive pediatric sarcomas.

## 2. MATERIALS AND METHODS

### 2.1. Cell culture and treatment

The following cell lines were used in this study: (i) a panel of in-house patient-derived OS (n=10), RMS (n=10) and ES (n=12) cell lines (listed in Supplementary Table 1) established from tumor tissues with written informed consent under previous projects (IGA MZCR NR9125-4, IGA MZCR NT13443-4, OP VVV CZ.02.1.01/0.0/0.0/16_019/0000868), which were approved for use in this study by the Masaryk University Research Ethics Committee, Brno, Czech Republic (approval no. EKV-2019-051); (ii) established sarcoma cell lines, namely, MNNG/HOS (OS, purchased from ECACC, #87070201), HuO-3N1 (OS, purchased from JCRB, #JCRB0413), SAOS-2 (OS, purchased from ATCC, #HTB-85) and RD (RMS, purchased from ECACC, #85111502); (iii) NTERA-2 (human pluripotent embryonal carcinoma cell line, clone D1) purchased from ECACC (#01071221), which served as a positive control; and (iv) the NDF-2 and NDF-3 cell lines (human neonatal dermal fibroblast cell lines, #CC-2509; Lonza Bioscience, Durham, NC, USA; a kind gift from Dr. Tomáš Bárta). All cell lines were authenticated by STR profiling (Generi Biotech, Hradec Králové, Czech Republic) and routinely tested for mycoplasma contamination via PCR. The culture media used are listed in Supplementary Table 2. All cell lines were cultured in a humidified 5% CO2 atmosphere at 37°C.

The following drugs targeting the p53 pathway were used for treatment: RO-5963 (MDM2/MDMX dual inhibitor, #444153, Sigma=:Aldrich, St. Louis, MO, USA) and PRIMA-1^MET^ (mut-p53 reactivator, #SML1789, Sigma=:Aldrich). Treatment with etoposide (#1226, TOCRIS Bio-Techne, Bristol, Great Britain) was used to stabilize p53 and activate its downstream signaling pathway in sarcoma cells. The drug treatments were performed the day after cell seeding.

### 2.2. Tumor samples

Formalin-fixed paraffin-embedded (FFPE) tumor tissue samples were obtained from 91 pediatric sarcoma patients (male/female: 54/37; age at diagnosis: 12 ± 5.6 years) treated at the University Hospital Brno or St. Anne’s University Hospital between 2006 and 2020. The study samples consisted of initial biopsy samples from 34 OS, 22 RMS (14 alveolar RMS, 7 embryonal RMS) and 36 ES patients. In total, 38 OS, 31 RMS and 41 ES biopsy samples were analyzed in this study. All samples were obtained with written informed consent, and their use in this study was approved by the Masaryk University Research Ethics Committee, Brno, Czech Republic (approval no. EKV-2019-051).

### 2.3. Western blotting

Western blotting and densitometry were performed as previously described [16]. The following antibodies were used for immunoblotting analysis: rabbit anti-SOX2 (1:1000; #3579, Cell Signaling Technology, CST, Danvers, MA, USA), rabbit anti-OCT4 (1:1000; #2750, CST), rabbit anti-NANOG (1:1000; #4903, CST), rabbit anti-c-MYC (1:1000; #5605, CST), rabbit anti-KLF4 (1:1000; #ab215036, Abcam, Cambridge, UK), mouse anti-p53 (1:10,000; #P6874, Sigma=:Aldrich), mouse anti-MDM2 (1:500; #sc-965, Santa Cruz Biotechnology, SCBT, Dallas, TX, USA), mouse anti-p63 (1:200, #sc-25268, SCBT), rabbit anti-p73 (1:1000, #sc-2750, SCBT), mouse anti-MDMX (1:200, #sc-74468, SCBT), rabbit anti-cleaved caspase-3 (1:1000; #9664, CST), rabbit anti-GAPDH (1:10,000, #2118, CST) and mouse anti-β-actin (1:10,000, #A5441, Sigma-Aldrich). GAPDH and β-actin served as loading controls.

### 2.4. Stemness index (SI) calculation

The protein expression levels of the stemness-associated transcription factors SOX2, OCT4, NANOG, KLF4 and c-MYC (relative to their appropriate loading controls) were used to calculate the SI for each sarcoma cell line. The SI of each cell line was determined as follows: sum of the expression values of individual stemness factors divided by the mean expression of the respective stemness factor in the given sarcoma subtype (OS, RMS, or ES).

### 2.5. RT□qPCR

RNA extraction, reverse transcription (with equal amounts of 20 ng of RNA per 1 µl of reaction mixture) and qPCR were performed as previously described [16]. The sequences of the primers used are listed in Supplementary Table 3. All qPCRs were performed in technical triplicates. To profile the expression of genes related to p53-mediated signal transduction based on the RT² Profiler™ PCR Array Human p53 Signaling Pathway (#330231, PAHS-027ZC, Qiagen, Hilden, Germany), reverse transcription and qPCR were performed using the RT^2^ First Strand Kit (#330401, Qiagen) and RT² SYBR Green ROX qPCR MasterMix (#330522, Qiagen), respectively, according to the manufacturer’s instructions.

### 2.6. *TP53* mutation analysis

In selected cell lines, amplicon ultradeep next-generation sequencing (NGS) of the whole-coding region of the *TP53* gene (exons 2–11 and canonical splice sites) was performed, and the results were assessed as previously described [17].

### 2.7. shRNA-mediated SOX2 knockdown

shRNA lentiviral particles were utilized to knock down the expression of SOX2 (#sc-38408-V, SCBT) according to the manufacturer’s instructions. Scrambled shRNA lentiviral particles (#sc-108080, SCBT) were used to establish corresponding controls. Single-cell clones were isolated from both stable shRNA control and SOX2-knockdown cell pools using puromycin (#sc-108071, SCBT) as the selection agent.

### 2.8. MTT assay

Cells were seeded in 96-well plates at densities of 5000 and 2000 cells/well and treated with the selected drugs for 72 h. After treatment, cell viability was measured using the MTT assay; thiazolyl blue tetrazolium bromide (#M2128, Sigma=:Aldrich) was added to a final concentration of 455 µg/ml. After 3 h of incubation under standard conditions, the medium was replaced with 200 μl of DMSO, and the absorbance was measured with a Sunrise absorbance reader (Tecan, Männedorf, Switzerland). The half-maximal inhibitory concentrations (IC_50_s) of the drugs were determined via nonlinear regression of the MTT assay data using GraphPad Prism 8.0.2 software (GraphPad Software, San Diego, CA, US), and the absolute IC_50_ values were determined according to the following formula: relative IC50*(((50-top)/(bottom-50))^(−1/hill slope)).

### 2.9. Limiting dilution sphere formation assay

The cells were harvested and dissociated into single-cell suspensions by adding Accutase (Biosera, Cholet, France); resuspended in serum-free, low-glucose DMEM (#LM-D1100, Biosera) supplemented with 10 ng/mL EGF (Sigma=:Aldrich), 20 ng/mL FGF2 (Sigma=:Aldrich), and 1× B-27 Supplement without vitamin A (Gibco, Carlsbad, CA, USA); and serially diluted and cultured as described previously [16]. Sphere formation was evaluated after one week, and only spheres ≥50 µm in diameter were counted using the IncuCyte® SX1 live cell imaging system (Sartorius, Göttingen, Germany). Frequencies of sphere-forming cells in the experimental groups were calculated and compared using ELDA software [18].

### 2.10. *In vivo* tumorigenicity assay and tissue microarray assembly

The cells were harvested and enzymatically dissociated, and a single-cell suspension of 1×10^6^ cells in 100 μl of low-glucose DMEM was injected subcutaneously into the right flank of nine-week-old NSG (NOD/ShiLtSz-*scid/I12rγ*^null^) mice. All animal experiments were conducted in accordance with a study (MSMT-4408/2016-6) approved by the Institutional Animal Care and Use Committee of Masaryk University and registered by the Ministry of Education, Youth and Sports of the Czech Republic as required by national legislation. The experiments were terminated after the mice reached a sufficient tumor volume (500 mm^3^) or after 25 weeks. Xenograft tissues were embedded in FFPE blocks and processed for tissue microarray (TMA) assembly. Briefly, tumor core biopsy samples (Ø3 mm) were punched from annotated donor blocks according to hematoxylin-eosin staining of the whole xenograft sections and re-embedded into a positionally encoded array of the recipient block cast manually using the Simport M473 T-Sue^TM^ Microarray Mold Kit (#M473-36, Simport Scientific, Beloeil, Canada) according to the manufacturer’s instructions.

### 2.11. Flow cytometry

After treatment, the cells were harvested and enzymatically dissociated into a single-cell suspension in PBS supplemented with 3% fetal bovine serum (#FB-1101, Biosera) and 2 mM EDTA (#ED2SS, Sigma=:Aldrich). After 15 minutes of incubation with 5 nM SYTOX™ Red Dead Cell Stain (#S34859, Invitrogen, Carlsbad, CA, USA) on ice, cell viability was assessed using a CytoFLEX S flow cytometer (Beckman Coulter, Brea, CA, USA).

### 2.12. Immunohistochemistry (IHC)

IHC staining was performed using a Ventana BenchMark ULTRA IHC/ISH System (Roche, Rotkreuz, Switzerland). For both the sarcoma xenograft TMA and human sarcoma tissue samples, 1.5-micron-thick sections were deparaffinized with EZ Prep Solution, antigen retrieval was performed with Cell Conditioning 1 reagent, and DAB staining was detected using the UltraView Universal DAB Detection Kit or the OptiView DAB IHC Detection Kit (all from Roche). The following antibodies were used: anti-SOX2 (#ab92494, Abcam), anti-OCT4 (#ab19857, Abcam), anti-NANOG (#ab109250, Abcam), anti-KLF4 (#ab215036, Abcam), anti-c-MYC (#ab32072, Abcam), anti-p53 (#05278775001, Roche), anti-MDM2 (#113-0230, Zytomed-Systems, Berlin, Germany), and anti-Ki-67 (*#*Ki681C01, DCS, Hamburg, Germany). The antibody dilutions, detection conditions and positive controls used are detailed in Supplementary Table 4.

The immunostained slides were evaluated by an experienced pathologist and scanned via a motorized widefield microscope (Zeiss AxioImager.Z2-ZEN) with a color camera (AxioCam 208) using ZEN Blue software (Zeiss, Praha, Czech Republic). QuPath software [19] was used to detect cell nuclei, classify tumor and nontumor cells (machine learning-based classifiers were trained by experienced pathologists) and perform DAB staining intensity analysis (machine learning-based intensity classifiers were trained on positive control tissues). The immunoreactivity of tumor cells was graded as negative (–), weak (+), medium (++), or strong (+++). For each of the detected proteins, histoscores were calculated as follows: 3 × percentage of cells with strong immunoreactivity + 2 × percentage of cells with medium immunoreactivity + 1 × percentage of cells with weak immunoreactivity.

### 2.13. Statistical analysis

All the statistical analyses were performed with at least three biological replicates (details in the figure legends) using GraphPad Prism 8.0.2 software. Each data point in the bar graphs represents an independent biological replicate. Unpaired one-/two-tailed Welch’s t-tests were performed for comparisons between two groups. Multiple-group comparisons were analyzed by one-way ANOVA with Welch’s correction followed by post hoc Dunnett’s test. The Spearman’s correlation coefficient was determined for correlation analyses. Kaplan=:Meier analyses were performed using the Mantel=:Cox test. Differences in the numbers of tumor spheres formed (assessed using ELDA software), and the numbers of metastases at diagnosis were compared via the chi-square test. p values < 0.05 were considered to indicate statistical significance; *p<0.05, **p<0.01, ***p<0.001.

## 3. RESULTS

### 3.1. SOX2 expression and p53 pathway dysregulation characterize tumorigenic stem-like sarcoma cells with aggressive growth features *in vivo*

To assess whether p53 pathway dysregulation is connected to the tumorigenic potential and stem-like traits of pediatric sarcomas, we first screened for the expression of canonical stemness-associated TFs—SOX2, OCT4, NANOG, KLF4, and c-MYC—together with p53 and MDM family proteins in a panel of 32 patient-derived and four established OS, RMS and ES cell lines (**Figs. 1a, S1**). Among these proteins, p63, p73 and MDMX showed no significant expression heterogeneity, and were thus excluded from further analyses (**Fig. S2**). Based on the expression of stemness-associated TFs, we calculated the stemness index (SI) of each sarcoma cell line (as detailed in Methods) and used it as a classifier potentially reflecting the stem-like capacity of sarcoma cells. We first identified SI^high^ (upper quartile SI) and SI^low^ (lower quartile SI) models in each subgroup of patient-derived cell lines and then aligned the established cell lines accordingly (**Fig. 1b**).

**Figure 1:**
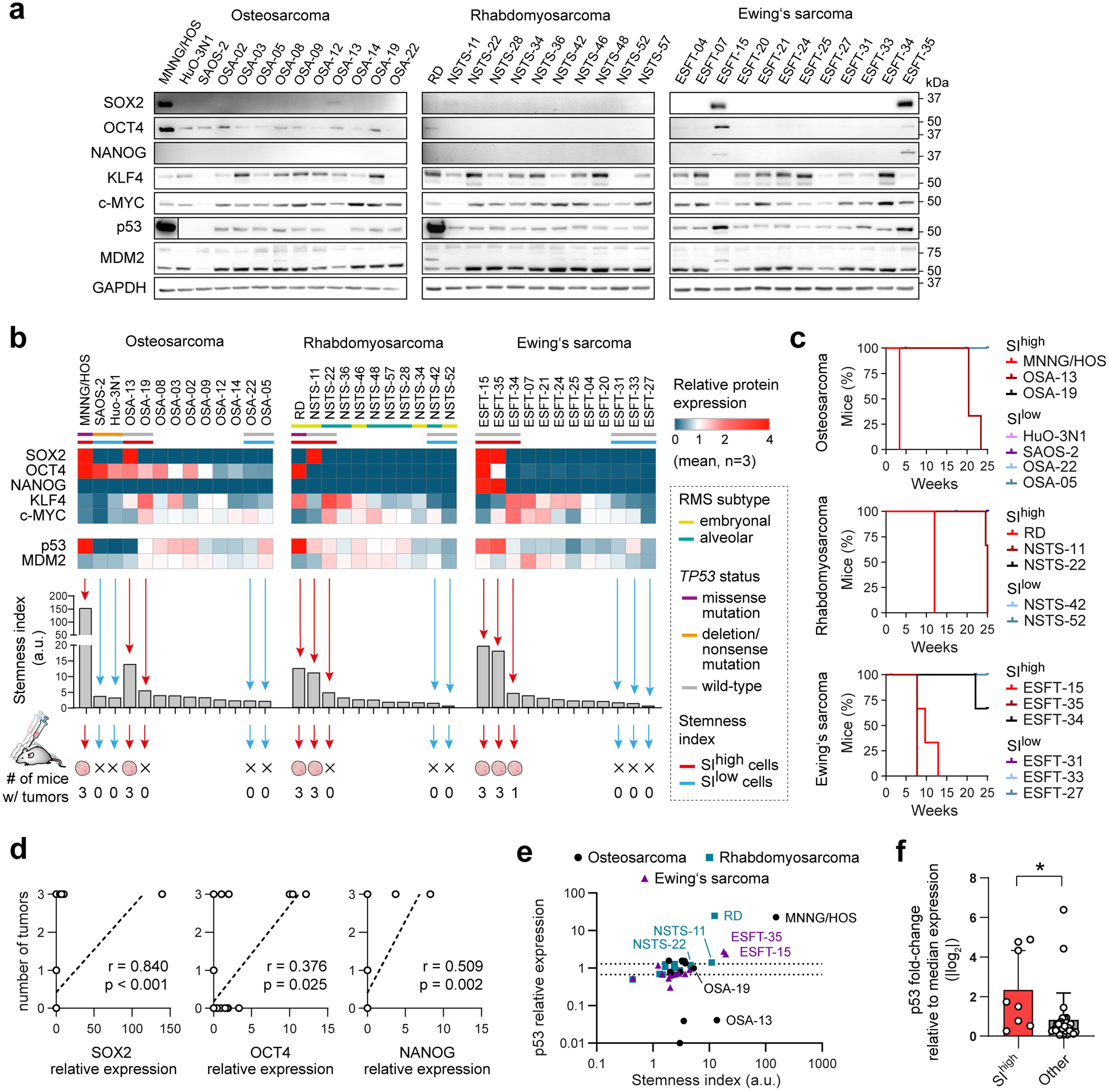
*In vitro* expression of the TFs SOX2, OCT4, and NANOG and p53 dysregulation are associated with pediatric sarcoma tumorigenicity. **a)** Representative images of western blot analysis (biological n=3) of the stemness-associated TFs SOX2, NANOG, OCT4, KLF4 and c-MYC and the p53 pathway proteins p53 and MDM2 in a panel of established and patient-derived OS, RMS and ES cell lines. The densitometric analysis is provided in Fig. S1. **b)** Heatmaps (top) of the mean protein expression levels (n=3) of the stemness-related TFs p53 and MDM2. The cell lines of each sarcoma subgroup were aligned according to the calculated SI (middle). The scheme (bottom) illustrates the tumorigenicity of the SI^high^ and SI^low^ cell lines assayed in NSG mice (n=3). **c)** Survival of xenograft model mice (n=3 mice per cell line) reflecting the tumor-initiating capacity and xenograft growth rates of the sarcoma cell lines tested. **d)** A significant positive correlation was found for the expression of the TFs SOX2, OCT4 and NANOG and the number of tumors formed by sarcoma cell lines in mice. **e)** Correlation between SIs and p53 protein levels in individual cell lines relative to the mean expression of each sarcoma subtype. Note that all tumorigenic SI^high^ models exhibit dysregulated p53 expression and fit within the upper or lower quartiles (indicated by dotted lines). **f)** The p53 levels in the SI^high^ models are significantly different from those in the other sarcoma cell lines. Data are presented as mean ± SD. Statistical significance was determined by one-tailed Welch’s t-test (**f**) and by Spearman’s correlation coefficient (**d**), *p<0.05, **p<0.01, ***p<0.001.

*In vivo* testing of SI^high^ and Sl^ow^ sarcoma models revealed SI as a good predictor of tumorigenicity in immunodeficient NSG mice, which is still considered the most important hallmark of cancer stemness [20]. Only the SI^high^ sarcoma cell lines (7/9) formed xenograft tumors in mice, whereas all SI^low^ cell lines (0/9 tested) were nontumorigenic (**Fig. 1b**). Notably, sarcoma models with the highest SIs rapidly initiated fast-growing tumors; the two SI^high^ cell lines that did not form tumors in mice had absolute SIs comparable to those of their nontumorigenic SI^low^ counterparts (**Fig. 1b, c**). These results suggested that the most differentially expressed stemness-associated TFs might be key factors related to the association between SI and tumorigenicity. Consistently, correlation analysis revealed that the number of tumors formed by the sarcoma cells was strongly correlated with the SOX2 level, and partially with OCT4 and NANOG (detected only in ES) levels (**Figs. 1d, S3**).

Importantly, the tumorigenic SI^high^ models showed either markedly increased or decreased p53 levels (relative to its median expression level across all cell lines) (**Fig. 1e**). A comparison of the SI^high^ models with the other cell lines revealed that p53 pathway dysregulation was significantly associated with the stemness of pediatric sarcoma cells (**Fig. 1f**). Interestingly, although the established MNNG/HOS and RD tumorigenic cell lines express mut-p53, no mutations in the coding region of the *TP53* gene were identified by NGS in the SI^high^ and SI^low^ patient-derived sarcoma cell lines (**Fig. 1b**), suggesting other mechanisms of p53 dysregulation. Collectively, these data indicated an association between cancer stemness and p53 dysregulation that might underly aggressive sarcoma cell phenotype.

We also asked whether the SIs established for patient-derived cell lines could serve as predictors of clinical outcomes. However, survival analysis revealed that the cell line-based SI, although informative of the CSC phenotype of the model itself, is unsuitable for patient outcome prediction and should thus be interpreted with caution (**Fig. S4**). Notably, the expression of all TFs and selected p53 pathway proteins was also evaluated at the mRNA level, but significantly less stable expression of individual transcripts was detected between biological replicates (**Fig. S5a**). Indeed, no association was found between the TF transcript levels and tumorigenicity (**Fig. S5b**), which underpins the importance of directly analyzing the expression levels of the encoded proteins to obtain proper insights into their roles in sarcomagenesis.

Tumor initiation in animal models involves the selection of cancer clones with the most malignant phenotype *in vivo* [21]. To analyze these phenotypes in pediatric sarcomas, xenografts obtained during the initial tumorigenicity assays were assembled into a self-constructed TMA and assessed via IHC (**Fig. 2a, d**). Following evaluation by an experienced pathologist, all slides were subjected to whole-slide scanning (**Figs. S6–9**) and computational image analysis using QuPath software, which allowed high-throughput quantitative expression analysis of individual sarcoma cells using machine learning classifiers (**Figs. 2b, S10a,** detailed results in **Figs. S11, S12**). This unbiased in-depth analysis revealed the same expression trends but provided higher sensitivity than manual readouts by a pathologist (**Figs. 2c, S10b,** detailed results in **Fig. S13**).

**Figure 2:**
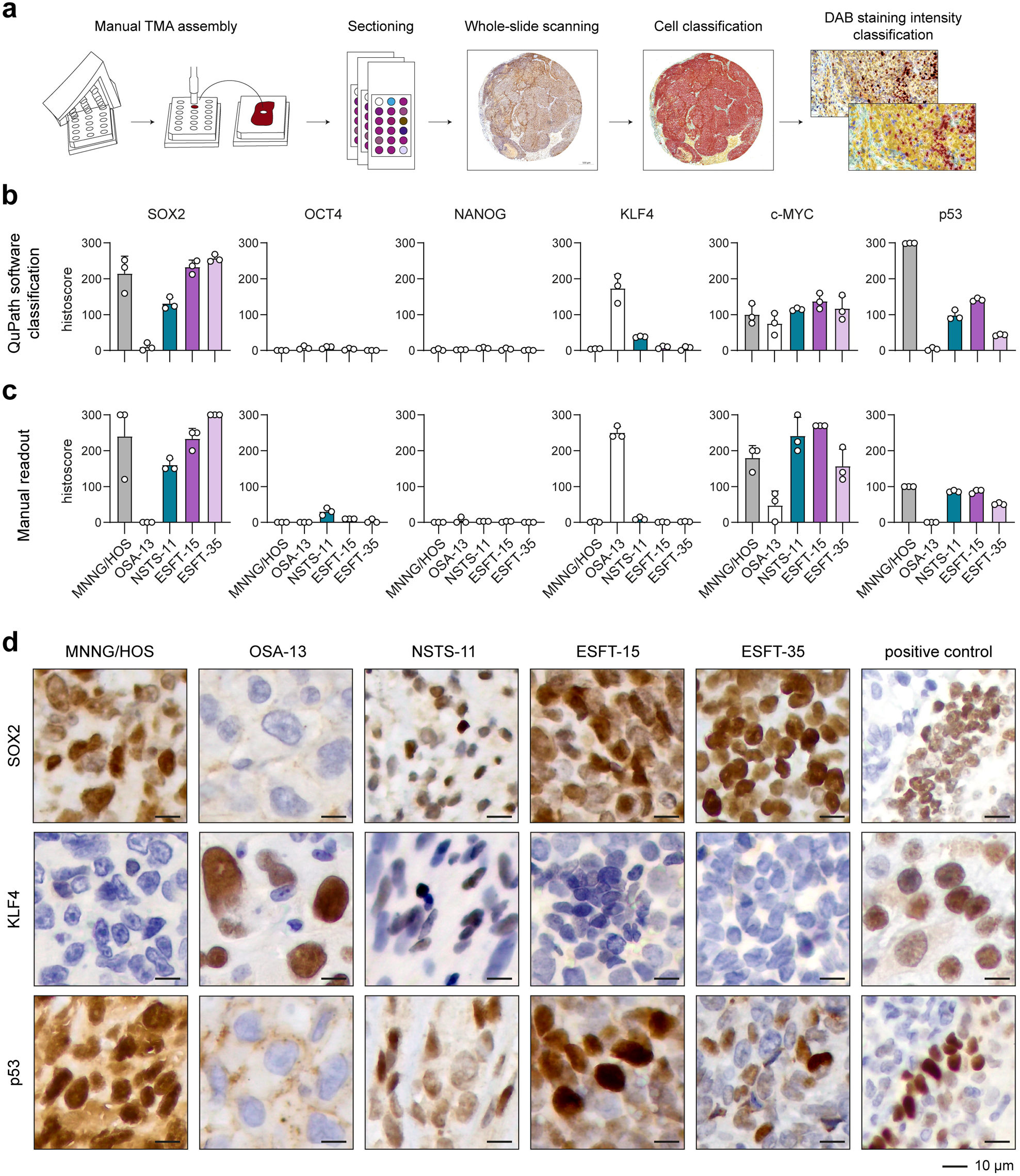
Cells with high SOX2 or KLF4 expression and dysregulated p53 expression are prevalent in sarcoma xenografts. **a)** Overview of the in-house pipeline for TMA assembly followed by IHC staining and machine learning-assisted image analysis of whole-slide scanned TMA sections. **b, c)** Histoscores, representing positivity and staining intensity of p53 and stemness-related TFs, assessed by computational analysis using QuPath software (**b**) or manual readout by an experienced pathologist (**c**). Data shown as mean ± SD, biological n=3. **d)** Representative images of IHC detection of SOX2, KLF4 and p53 in xenograft tumors formed by the indicated cell lines or in positive control tissues (SOX2, fetal lungs; KLF4, seminoma; p53, tonsilla).

IHC revealed that most sarcoma xenografts maintained high expression of SOX2 but generally lacked OCT4 and NANOG expression, indicating the crucial role of SOX2 in sarcomagenesis (**Fig. 2b, c**). Moreover, abundant expression or complete absence of p53 was observed in all analyzed sarcoma xenograft tissues (**Fig. 2b, c**). An interesting phenotypic switch and KLF4 upregulation, likely compensating for loss of SOX2 expression, was observed in the OS model OSA-13, negative for p53 protein expression.

### 3.2. SOX2 knockdown reduces the sarcosphere formation capacity in the context of mut-p53

Our results revealed that the stemness-associated TF SOX2 and p53 pathway dysregulation are associated with the CSC phenotype of pediatric sarcoma cells. We therefore sought to elucidate whether different p53 backgrounds (mut-p53 vs. wt-p53) affect how SOX2 contributes to sarcoma stemness. We thus established single-cell clones with stable shRNA-mediated knockdown of SOX2 (shSOX2) derived from mut-p53 MNNG/HOS and wt-p53 ESFT-15 cells (**Fig. 3a, b**). Immunoblotting revealed that viable clones of MNNG/HOS and ESFT-15 cells with SOX2 knockdown exhibited marked upregulation of NANOG and KLF4 expression, respectively (**Figs. 3a, S14a**), indicating possible compensation for self-renewal in the isolated single-cell clones. These findings agree with the phenotypic switch observed in sarcoma xenografts, where KLF4 was likely compensating p53 loss to maintain the tumor-initiating capacity of OS patient-derived OSA-13 cells (**Fig. 2b**). Upregulation of *KLF4* expression in ESFT-15-shSOX2 cells was also detected at the mRNA level (**Fig. 3b**), together with increase in *OCT4* and *NANOG* and decrease in *MYC* transcripts, although these changes did not lead to significant changes in the corresponding protein levels (**Figs. 3a, b, S14a, b**).

**Figure 3:**
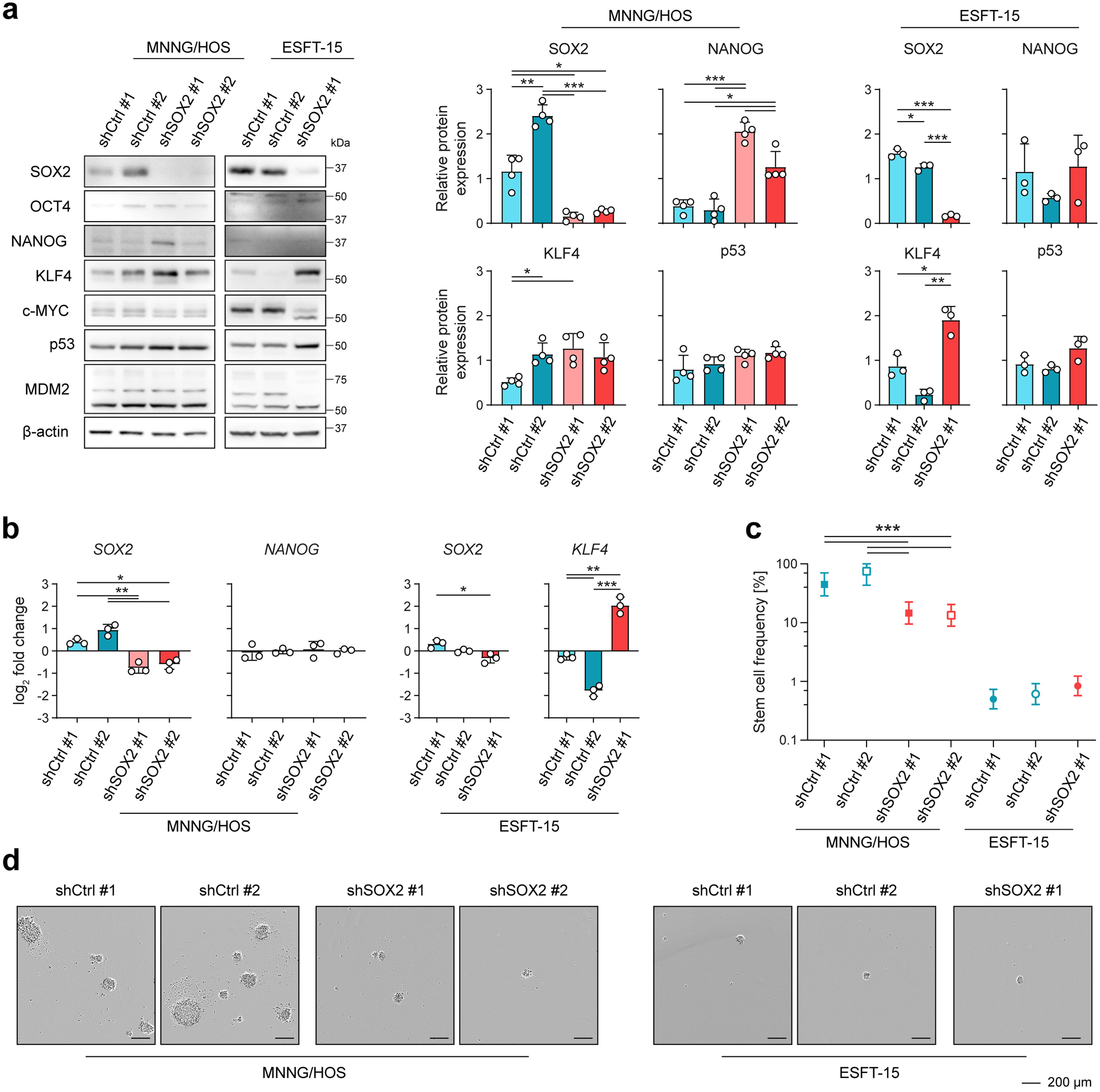
Knockdown of SOX2 expression is associated with suppression of sphere-formation capacity in sarcoma cells harboring mut-p53. **a)** Representative western blot images (left) and densitometric analysis (right) of stemness-associated TFs, p53 and MDM2 in mut-p53 MNNG/HOS and wt-p53 ESFT-15 clones with shRNA-mediated knockdown of SOX2 (shSOX2) compared with their scramble shRNA controls (shCtrl). Data presented as mean ± SD, biological n=3–4. Densitometric analysis of the OCT4, c-MYC, and MDM2 proteins is provided in Fig. S10a. **b)** qPCR analysis of selected stemness-associated TFs in MNNG/HOS and ESFT-15 shSOX2 clones. Data are presented as mean ± SD, biological n=3, technical n=3. Analysis of remaining genes is provided in Fig. S10b. **c)** Sphere formation assay evaluating the stem-like phenotype of MNNG/HOS and ESFT-15 sarcoma cells after SOX2 knockdown. The stem cell frequencies and probabilities were computed using ELDA software [18]. Data are shown as mean ± 95% confidence interval, biological n=3–4, technical n=5. **d)** Representative images of sarcospheres formed by shCtrl and shSOX2 clones of the MNNG/HOS and ESFT-15 cell lines. All experiments were performed using single cell-derived clones (indicated by numbers). Statistical significance was determined by one-way ANOVA with Welch’s correction followed by post hoc Dunnett’s test (**a, b**) or chi-square pairwise test (**c**), *p<0.05, **p<0.01, ***p<0.001.

To assess whether SOX2 expression downregulation affects the stemness of sarcoma cells in a p53 status-dependent manner, both shSOX2 and shRNA control (shCtrl) clones were subjected to a sphere formation assay, which is the most comprehensive functional assay for evaluating the CSC phenotype *in vitro*. Unlike wt-p53 ESFT-15-shSOX2 cells, MNNG/HOS-shSOX2 cell clones carrying mut-p53 showed significantly reduced sphere-formation capacity compared with their shCtrl counterparts (**Fig. 3c, d**). Consistent with the later tumor onset observed *in vivo* (**Fig. 1c**), wt-p53 ESFT-15-shCtrl cells formed markedly fewer spheres than MNNG/HOS-shCtrl clones with mut-p53 (**Fig. 3c**). These results indicate that *TP53* mutation is important in the development of the highly aggressive CSC phenotype in pediatric sarcoma and further synergizes with the increased level of the stemness-associated TF SOX2.

### 3.3. Restoring p53 activity selectively induces cell death in aggressive stem-like Ewing’s sarcoma cells

Because p53 dysregulation was associated with sarcoma stemness, we next explored whether p53 reactivation might selectively target sarcoma CSCs and impair their viability. Given the wt-p53 status of the patient-derived cell lines in this study, the effects of two compounds with different mechanisms of action—a dual inhibitor of MDM2/MDMX, RO-5963 [22], and a control mut-p53 reactivator, PRIMA-1^MET^ [23]—were assessed in paired SI^high^ and SI^low^ cell lines from each of the sarcoma subgroups. Among the SI^high^ and SI^low^ pairs, differential sensitivity to p53 reactivation was observed only in the ES pair (**Fig. 4a**). When exposed to p53-targeted therapy, SI^high^ ESFT-15 cells were responsive to ∼40-fold lower concentrations of RO-5963 (absolute IC_50_ values of 1.5 µM vs. 56.4 µM) than SI^low^ ESFT-27 cells (**Fig. 4a**). Unsurprisingly, compared with the effects in healthy human fibroblasts (**Fig. 4b**), treatment with PRIMA-1^MET^, which targets mut-p53, did not exert selective effects on the ES pair (**Fig. 4a**). Exploring the vulnerability of SI^high^ ES cells to MDM2/MDMX inhibition revealed that ESFT-15 cells exhibited markedly elevated p53 expression coupled with compensating features such as dysregulation of p53 target genes involved in cell cycle arrest, DNA repair, and apoptosis (**Figs. 4e, f, S15, S16**). Interestingly, the p53 pathway remained responsive in ESFT-15 cells, and these cells exhibited the highest increase in p53 activity upon treatment-induced p53 stabilization (**Figs. 4e, f, S15, S16**), suggesting that MDM2/MDMX inhibitors might be effective even against tumors with upregulated wt-p53 expression.

**Figure 4:**
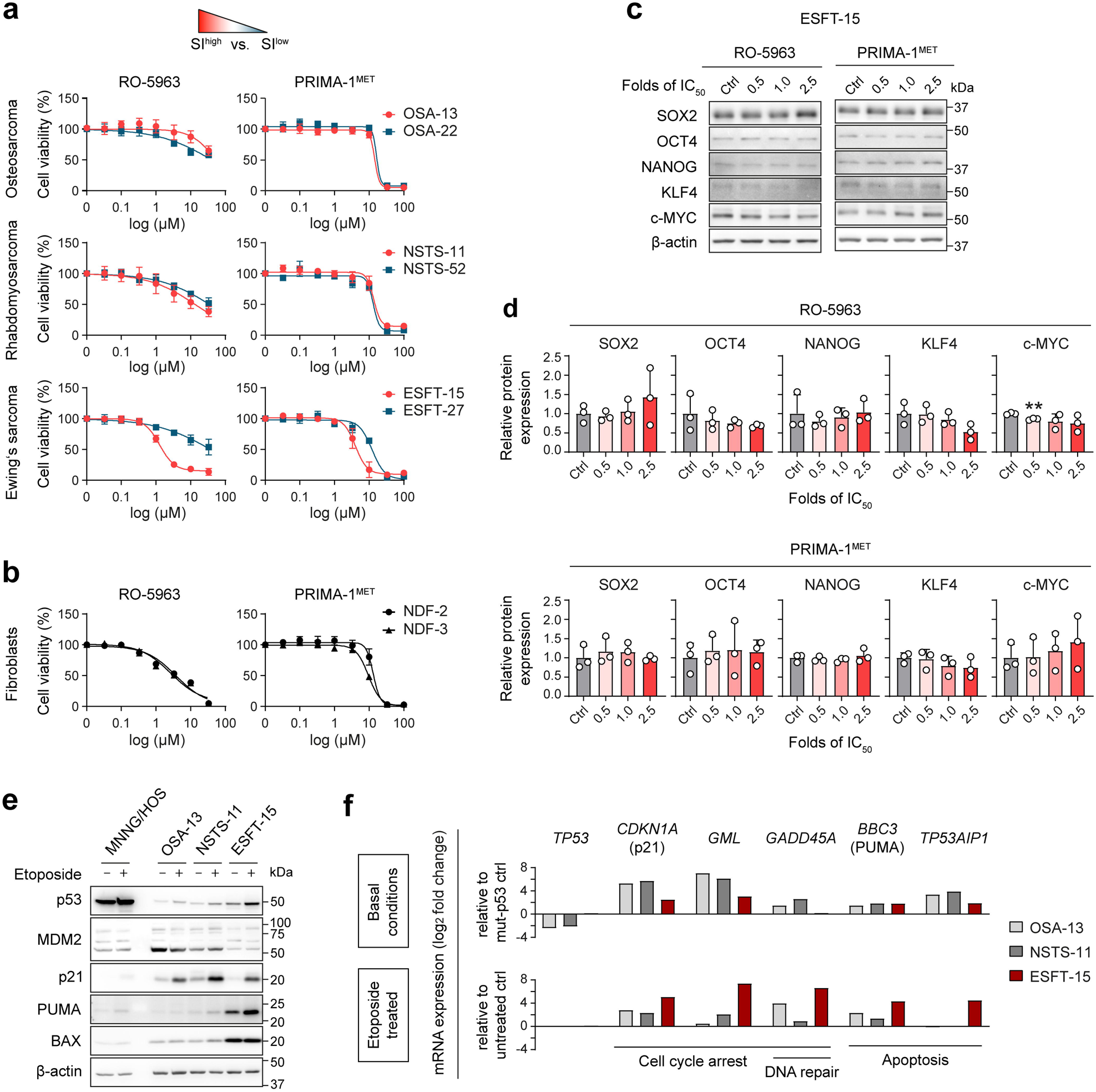
Stem-like Ewing’s sarcoma cells are vulnerable to p53 reactivation. **a)** MTT cell viability assay of the SI^high^ and SI^low^ OS, RMS and ES cell lines after 72 h of treatment with RO-5963 or PRIMA-1^MET^. Notably, only cells derived from ES showed marked differences in sensitivity to p53 reactivation. Data are presented as mean ± SD, biological n=3–5, technical n=3–5. **b)** MTT cell viability assay of control healthy fibroblast cell lines after 72 h of exposure to RO-5963 or PRIMA-1^MET^. Data are presented as mean ± SD, biological n=3, technical n=3–5. **c, d)** Representative western blot images (**c**) and densitometric analysis (**d**) of stemness-associated TFs expression in aggressive ESFT-15 cells after 72 h of treatment with RO-5963 or PRIMA-1^MET^. Ctrl, DMSO-treated control. Data shown as mean ± SD, biological n=3. **e)** Representative western blot images (n=3) showing the expression of p53 and its downstream targets in the mut-p53 (MNNG/HOS) and wt-p53 SI^high^ cell lines after 24 h of treatment with 25 µM etoposide, which induces DNA damage. **f)** qPCR analysis of the mRNA expression of *TP53* and its target genes regulating cell cycle arrest, DNA repair and apoptosis: upper panel, basal levels relative to those in the control (ctrl) mut-p53 MNNG/HOS cells; lower panel, levels in cells after 24 h of treatment with 25 µM etoposide relative to those in the respective untreated controls (ctrl); biological n=1. Statistical significance was determined by one-way ANOVA with Welch’s correction followed by post hoc Dunnett’s test (**d**), **p<0.01.

To elucidate how p53 reactivation affects sarcoma stemness, we analyzed the effects of RO-5963 on sensitive tumorigenic ESFT-15 cells. Interestingly, similar to PRIMA-1^MET^, RO-5963 treatment did not significantly alter the expression of any of the examined stemness-associated TFs (**Fig. 4c, d**), suggesting that the selective antiproliferative effects of RO-5963 observed in SI^high^ ES cells are unlikely to be mediated by induction of differentiation. To investigate the underlying mechanism, we analyzed the expression of p53, MDM2 and cleaved caspase-3 in ESFT-15 cells and control healthy fibroblasts treated with increasing concentrations (folds of IC_50_) of the drugs (**Fig. 5a, b**). Unlike PRIMA-1^MET^ treatment, inhibition of the MDM2/MDMX-p53 interaction by RO-5963 led to p53 upregulation and induced apoptosis in ESFT-15 cells, as indicated by a dose-dependent increase in the cleaved caspase-3 levels (**Fig. 5a, b**). At least partial upregulation of p53 was also detected in both fibroblast cell lines; however, the induction of cleaved caspase-3 was either not detected or limited. Indeed, flow cytometry confirmed that RO-5963 treatment markedly induced cell death in SI^high^ ES cells, whereas its cytotoxic effects on healthy fibroblasts were not dose-dependent and remained limited even at the highest drug concentration (58.3 ± 2.6% vs. 4.8 ± 0.3% and 7.5 ± 1.9% for ESFT-15 vs. NDF-3 and NDF-2; **Fig. 5c, d**).

**Figure 5:**
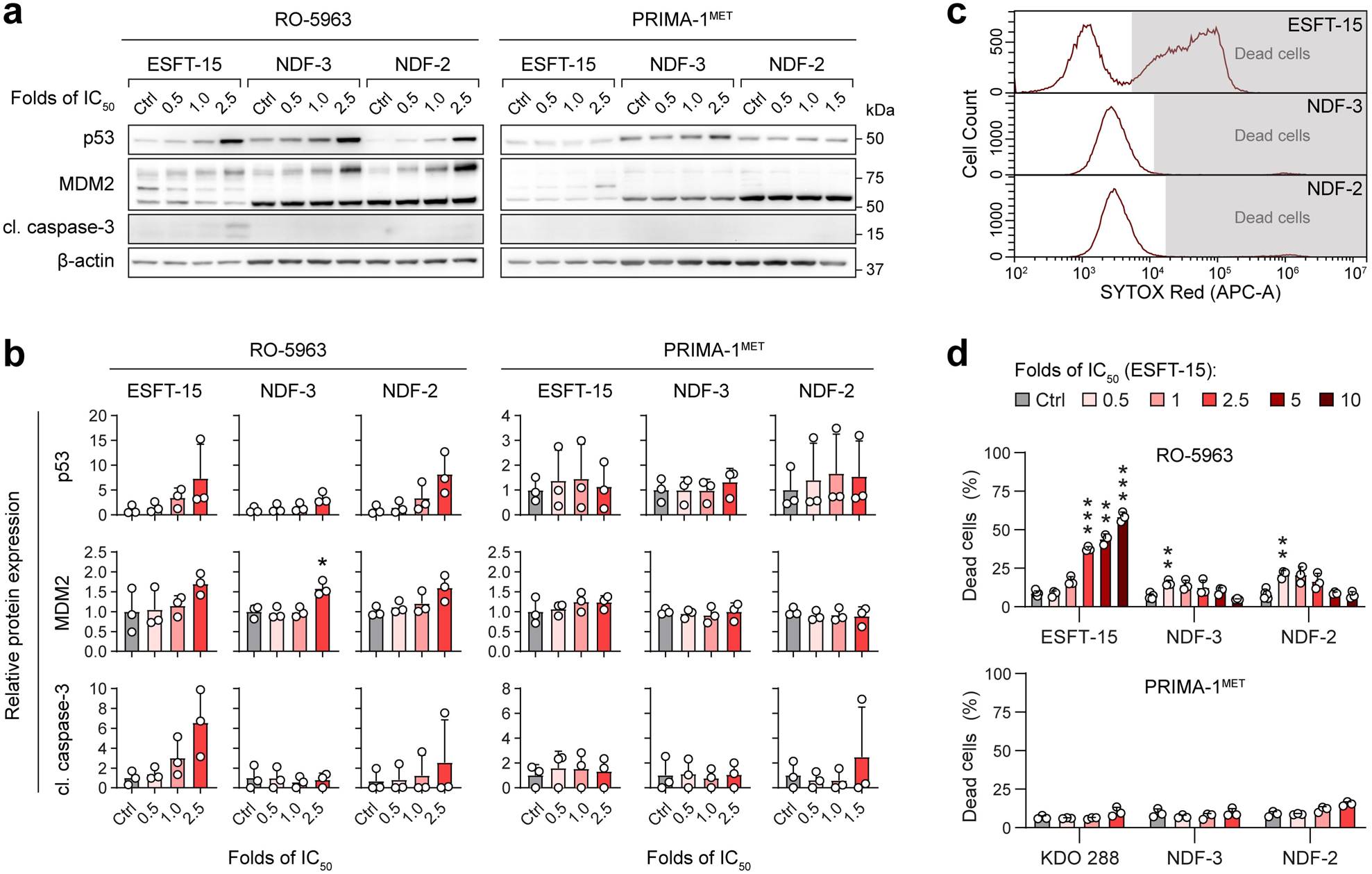
Inhibition of the MDM2/MDMX-p53 interaction potently activates the p53 pathway and induces cell death in aggressive Ewing’s sarcoma cells without marked cytotoxicity in healthy fibroblasts. **a, b)** Representative western blot images (**a**) and densitometric analysis (**b**) of p53, MDM2 and cleaved caspase-3 expression in the ESFT-15 ES cell line and healthy fibroblasts NDF-3 and NDF-2 after 72 h of exposure to RO-5963 and PRIMA-1^MET^. Data are presented as mean ± SD, biological n=3. **c)** Representative flow cytometry histograms showing marked differences in SYTOX Red-positive dead cells in the ESFT-15 and fibroblast cell lines after 72 h of exposure to the 10×IC_50_ dose (ESFT-15) of RO-5963. **d)** Flow cytometry analysis of dead vs. live cells after 72 h of treatment with the p53 pathway-targeting drugs RO-5963 and PRIMA-1^MET^. Data presented as mean ± SD, biological n=3-6. Ctrl, DMSO-treated control. Statistical significance was determined by one-way ANOVA with Welch’s correction followed by post hoc Dunnett’s test (**b, d**), *p<0.05, **p<0.01, ***p<0.001.

### 3.4. p53 associates with SOX2 expression in ES and is frequently expressed in initially metastatic sarcomas

To assess the clinical relevance of the identified link between the stemness expression signature and p53 pathway dysregulation, we performed IHC on tumor tissues from 91 pediatric sarcoma patients. A pathologist assessed 38 OS, 31 RMS and 41 ES tumor samples for the expression of p53, SOX2 and KLF4, which played crucial roles in sarcoma initiation in our models. Interestingly, this analysis revealed predominant expression of only one of the selected stemness-associated TFs in OS and ES. While KLF4 was almost exclusively expressed in OS tissues (28/38), SOX2 was not detected in any of the OS samples (**Fig. 6a**). Conversely, upregulation of SOX2 but not KLF4 expression was prevalent in ES tissues (12/41 vs. 1/41) (**Fig. 6a**). SOX2 (4/31) and KLF4 (10/31) positivity without marked overexpression of one of these factors was observed in RMS samples (**Fig. 6a**).

**Fig 6:**
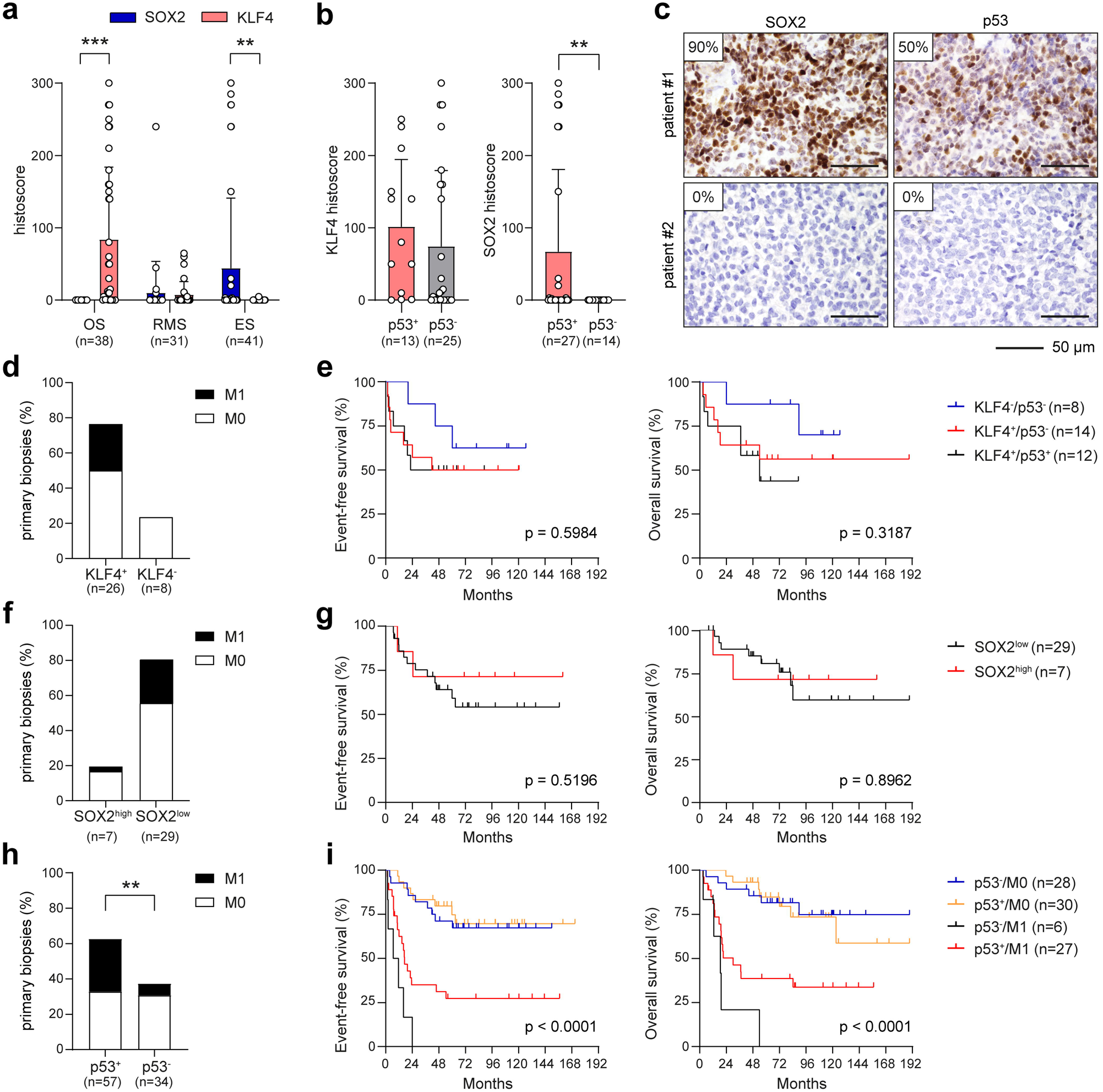
p53 positivity associates with SOX2 expression in ES and is more prevalent in sarcomas with metastasis at diagnosis. **a)** IHC analysis of SOX2 and KLF4 expression in OS, RMS and ES biopsies. **b)** Expression of KLF4 and SOX2 in p53-positive and p53-negative OS and ES tissues, respectively. In ES, SOX2 was detected only in p53-positive biopsies. **c)** Representative images of IHC analysis for SOX2 and p53 in ES tissue. **d)** Frequency of metastases at diagnosis in OS patients with KLF4-positive and KLF4-negative tumors. **e)** Kaplan=:Meier analysis of OS patients stratified by KLF4 and p53 protein expression. **f)** Frequency of metastases at diagnosis in ES patients with SOX2^high^ and SOX2^low^ tumors. **g)** Kaplan=:Meier analysis of patients with ES stratified by SOX2 protein expression (the cutoff for a SOX2^low^ histoscore was ≤ 45). **h)** Frequency of metastases at diagnosis in pediatric sarcoma patients with p53-positive and p53-negative tumors. The results show that p53 expression is associated with early metastatic disease. **i)** Kaplan=:Meier analysis of sarcoma patients stratified by p53 protein expression and the presence of initial metastases. M0, no metastatic events; M1, presence of initial metastases. Statistical significance was determined by Welch’s t-test (**a, b**), the chi-square test (**d, f, h**) and the Mantel–Cox test (**e, g, i**), *p<0.05, **p<0.01, ***p<0.001.

Correlation analysis revealed that KLF4 was not associated with p53 positivity in OS samples (**Fig. 6b**). However, SOX2 was detected only in p53-overexpressing ES tissues (**Fig. 6b, c**), further supporting our findings that p53 dysregulation is an important promoter of sarcoma stemness. To address whether SOX2, KLF4 and/or dysregulated p53 are clinically useful prognostic biomarkers in our pediatric sarcoma cohort, the correlations of their expression with clinicopathological features and survival were analyzed using primary biopsy samples (n=91). This revealed that in the OS cohort, only patients with KLF4-positive tumors present with metastases at diagnosis (**Fig. 6d**). Moreover, regardless of the p53 level, KLF4 positivity tended to be associated with worse survival in OS patients (**Fig. 6e**). High SOX2 expression was recently suggested as an independent biomarker of poor prognosis in ES patients [24]. To analyze the prognostic significance of SOX2 in our ES cohort, we therefore applied a matching histoscore-based dichotomization: SOX2^low^ (histoscore ≤45) and SOX2^high^ (histoscore >45). Strikingly, this analysis did not reveal any correlation between high SOX2 expression and the presence of metastases at diagnosis (**Fig. 6f**) or a significant difference in the ES patient survival rate (**Fig. 6g**).

In contrast, p53 positivity was significantly more prevalent in sarcoma patients with early metastases (**Fig. 6h**). Intriguingly, a trend toward improved overall survival (p = 0.09) was observed in p53-positive metastatic sarcomas (**Fig. 6i**), suggesting that the absence of p53 expression in the context of already metastatic disease might be associated with the most unfavorable prognosis.

Collectively, our results indicate that the KLF4, SOX2 and p53 protein levels assessed via IHC have rather limited value as independent biomarkers of clinical outcomes/prognosis for pediatric sarcomas. However, p53-positive staining is significantly associated with metastatic disease, which supports our experimental findings and indicates that restoring wt-p53 function is a promising strategy for the clinical treatment of aggressive sarcomas.

## 4. DISCUSSION

Pediatric sarcomas are highly malignant tumors with unmet clinical needs. We and others have provided evidence linking the aggressive behavior of sarcoma cells to the CSC phenotype [9,10,25–27]. However, the primary determinants of sarcoma stemness and, more crucially, their clinical relevance remain poorly understood. In this study, we utilized an extensive panel of pediatric sarcoma models and tissue samples to explore whether dysregulation of p53, a recently identified regulator of stemness [1,28], is connected with the stem-like phenotype of sarcoma cells. We showed that altered p53 function is associated with sarcoma stemness and aggressiveness and might therefore be exploited for developing novel therapies for at least some pediatric sarcomas that are refractory to available treatments.

Although numerous markers of sarcoma stemness have been suggested [3,5–7], recent findings have emphasized that future studies should focus on investigating the role of core stemness-associated TFs [9]. Indeed, the newly introduced SI, which reflects differential expression of stemness-associated TFs, was found to be a valuable estimate of cancer stemness, predicting the tumorigenicity of sarcoma cells in this study. Our initial screening highlighted SOX2 as a major marker of sarcoma stemness and showed that only cells expressing SOX2 or KLF4 were prevalent in sarcoma xenografts. The expression of OCT4 and/or NANOG detected in sarcoma cells *in vitro* was markedly suppressed or absent *in vivo*, which contradicts the previously suggested role of these TFs in sarcoma tumorigenicity [6,7,27]. Notably, our observation that KLF4 expression was almost exclusive to OS xenografts lacking SOX2 was further corroborated by IHC of human OS tissues. Consistent with experimental studies indicating that KLF4 is a promoter of stemness and metastasis in OS [13,29–31], we detected abundant KLF4 positivity associated with a trend toward worse survival in our OS cohort, suggesting that KLF4 rather than SOX2 is a clinically relevant oncoprotein in OS. In contrast, SOX2 expression was predominant in ES tumors and exclusively expressed in p53-positive patients. SOX2 was recently identified as an independent marker of poor prognosis in patients with ES [24] and sarcomas in general [26]. However, our results question its prognostic utility, as we did not find any association between SOX2 expression and ES patient survival. The high KLF4 and SOX2 expression in the respective sarcoma subtypes agrees with previous studies [31,32].

Altered p53 expression has been suggested to be an important promoter of sarcomagenesis in rodents and mesenchymal stem cell-derived models [33–36]. Consistently, data obtained from a large panel of pediatric sarcoma cell lines revealed that a high SI and enhanced sarcoma tumorigenicity are associated with *TP53* mutations and other p53 pathway alterations. This finding was substantiated by the high p53 positivity rate in sarcoma tissues of patients with early metastatic disease. However, p53 expression failed to discriminate pediatric sarcoma patients with poor outcomes, highlighting another contradictory result between our study and previous studies investigating the prognostic value of p53 in sarcomas [37–41]. Interestingly, none of our SI^high^ or SI^low^ patient-derived sarcoma cell lines carried *TP53* mutations, which disagrees with previously reported p53 mutation rates, particularly for OS [1]. Nonetheless, we showed that wt-p53 SI^high^ ES cells, characterized by elevated basal expression of p53 but dysregulated levels of its downstream targets, are highly vulnerable to additional pharmacological stabilization of p53, leading to hyperactivation of p53 signal transduction. These results demonstrate that simple analysis of the *TP53* DNA sequence is insufficient for assessing the p53 status and that evaluating the p53 protein levels might help identify patients with dysregulated p53 signaling who would benefit from combined regimens involving p53-stabilizing therapies. This is particularly important because wt-p53 activity may be suppressed by various protein=:protein interactions, including not only well-known interactions of p53 with MDM molecules [42] but also with specific oncoproteins, such as EWS-FLI-1 that drives ES development [43].

Under normal conditions, p53 is a crucial factor regulating the reprogramming and maintenance of stemness in induced pluripotent stem cells [44]. Consistently, p53 dysregulation was associated with the expression of the core stemness factor SOX2 in ES tissues. Strikingly, SOX2 depletion decreased but did not entirely suppress the stemness of mut-p53 sarcoma cells, suggesting that p53 alteration is important for acquisition of a fully stem-like phenotype in sarcomas. Moreover, SOX2 knockdown resulted in upregulated expression of NANOG and KLF4 (both associated with cancer stemness [45,46]) in the MNNG/HOS and ESFT-15 cell lines, respectively. These outcomes are inconsistent with prior studies, which suggested that SOX2 downregulation reduces NANOG expression [47] and KLF4 protein stability [48].

Because dysregulated p53 was linked to an aggressive stem-like sarcoma phenotype, we explored p53 reactivation as an approach to target sarcoma CSCs. We leveraged our paired OS, RMS and ES models with significant differences in SIs and thus tumorigenic potential to compare the effects of p53 pathway-targeting agents. Using this approach, we demonstrated that SI^high^ tumorigenic ES cells, but not their SI^low^ counterparts, were highly sensitive to p53 reactivation. Inhibiting the MDM2/MDMX-p53 interaction effectively induced p53 expression and apoptosis in stem-like ES cells harboring wt-p53 (present in most ES cases [1]) but showing basal dysregulation of p53 downstream effectors. RO-5963 increased the p53 protein levels at least partially in both ESFT-15 and fibroblast cells but markedly promoted cell death only in the tumorigenic SI^high^ cell line without significant effect on healthy fibroblasts, suggesting a potentially favorable therapeutic window. Consistently, RO-5963 previously showed similar potential for p53-targeted therapy in OS, as it induced apoptosis in cells with intact *TP53* [22].

In conclusion, our findings strongly suggest a significant association among p53 pathway dysregulation, the enhanced stemness-related transcription network (marked by SOX2 or KLF4 expression) and the aggressiveness of sarcoma cells. Clinically, alterations in p53 expression were prevalent in metastatic sarcomas, providing a relevant target for reactivation therapies. We demonstrated that inducing p53 activity via MDM2/MDMX inhibition selectively targeted tumorigenic stem-like ES cells characterized by high basal levels of wt-p53 but had no significant effect on healthy fibroblasts. Although further preclinical studies are needed to validate these results, p53 reactivation might be a promising strategy for treating refractory/metastatic p53-overexpressing pediatric sarcomas, regardless of their *TP53* mutational status.

## ABBREVIATIONS

CSC: cancer stem cell
ES: Ewing’s sarcoma
FFPE: formalin-fixed paraffin-embedded
IHC: immunohistochemistry
mut-p53: mutant p53
NGS: next-generation sequencing
OS: osteosarcoma
RMS: rhabdomyosarcoma
shCtrl: shRNA control
shSOX2: shRNA-mediated knockdown of SOX2
SI: stemness index
TF: transcription factor
TMA: tissue microarray
wt-p53: wild-type p53

## ETHICS STATEMENT

This study was conducted in accordance with the Declaration of Helsinki and approved by the Research Ethics Committee of Masaryk University (approval no. EKV-2019-051; grant on April 14, 2020). All human tumor tissues and noncommercial patient-derived sarcoma cell lines were obtained with written informed consent from patients or their legal guardians. The animal studies were approved by the Institutional Animal Care and Use Committee of Masaryk University and were registered by the Ministry of Education, Youth and Sports of the Czech Republic (ref. no. MSMT-4408/2016-6) as required by national legislation.

## CREDIT AUTHORSHIP CONTRIBUTION STATEMENT

**Lucie Curylova**: Writing – original draft, Investigation, Methodology, Visualization, Data curation, Formal analysis, Writing – review & editing. **Iva Staniczkova Zambo**: Investigation, Visualization, Writing – review & editing. **Jakub Neradil**: Investigation, Writing – review & editing. **Michal Kyr**: Data curation, Formal analysis, Writing – review & editing. **Nicola Jurackova**: Methodology, Investigation. **Sarka Pavlova**: Investigation, Writing – review & editing. **Kristyna Polaskova**: Data curation. **Peter Mudry**: Resources. **Jaroslav Sterba**: Resources. **Renata Veselska**: Resources, Funding acquisition. **Jan Skoda**: Conceptualization, Writing – original draft, Formal analysis, Visualization, Funding acquisition, Supervision, Project administration, Writing – review & editing. All authors read and approved the final version of the manuscript.

## ACKNOWLEDGEMENTS

LC was supported by the Brno Ph.D. Talent Scholarship funded by the Brno City Municipality. JN, KP, RV and JS (Jan Skoda) acknowledge the project National Institute for Cancer Research (Programme EXCELES, ID Project No. LX22NPO5102)—Funded by the European Union—Next Generation EU. SP acknowledges the conceptual development of the research organization (FNBr 65269705) provided by the Ministry of Health of the Czech Republic. The authors thank Dr. Jan Verner and Johana Marešová for their skillful technical assistance and Dr. Tomáš Bárta (Department of Histology and Embryology, Faculty of Medicine, Masaryk University), for providing the normal fibroblast lines.

