## Supplementary Information for "Dysregulation of the p53 pathway provides a therapeutic target in aggressive pediatric sarcomas with stem-like traits"

### This PDF file includes:

Figures S1 to S16

Supplementary Tables 1 to 4

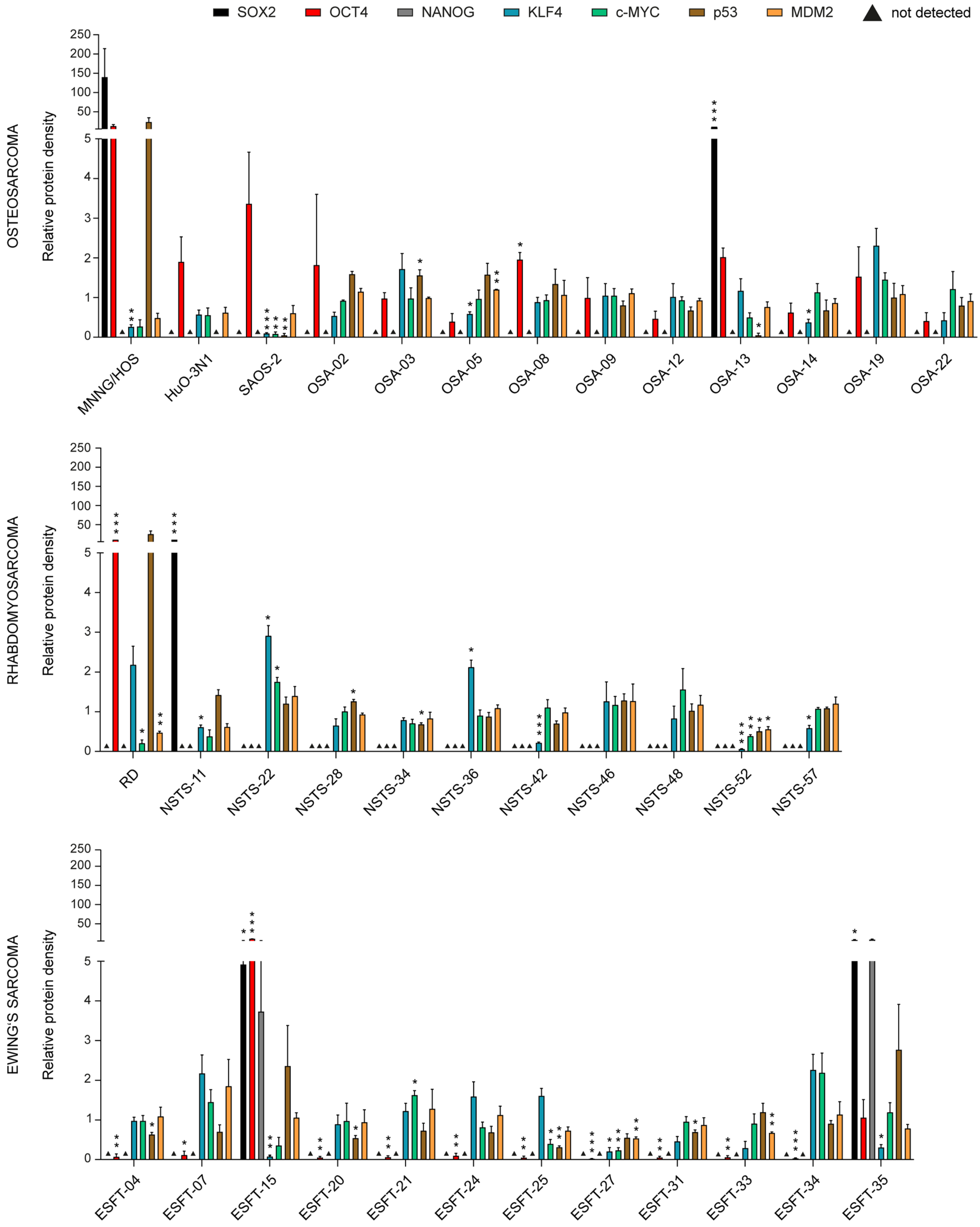


**Figure S1: Densitometric analysis of the protein expression of selected stemness-associated TFs, p53 and MDM2 in a panel of OS, RMS and ES established and patient-derived cell lines.** Expression values of each cell line were evaluated relative to the average expression value of the respective protein in each subgroup of patient-derived sarcomas. Data are mean ± SD, biological n=3. Statistical significance was determined by one-way ANOVA with Welch's correction followed by post-hoc Dunnett's test, *p<0.05, **p<0.01, ***p<0.001.

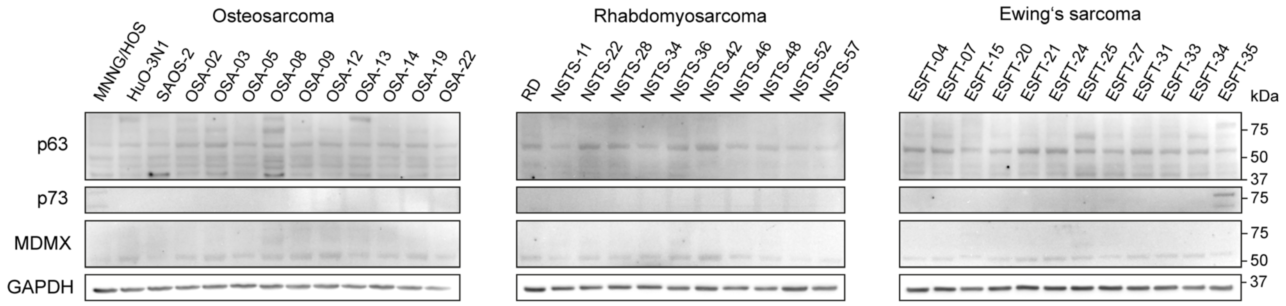


**Figure S2: Representative western blot images of p63, p73 and MDMX expression in sarcoma cell lines.** Biological n=3.

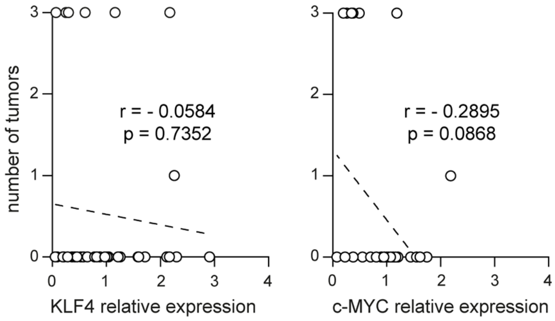


**Figure S3: Correlation analysis between number of tumors formed by sarcoma cell lines in mice and expression of KLF4 and c-MYC TFs.** r was determined by Spearman correlation test.

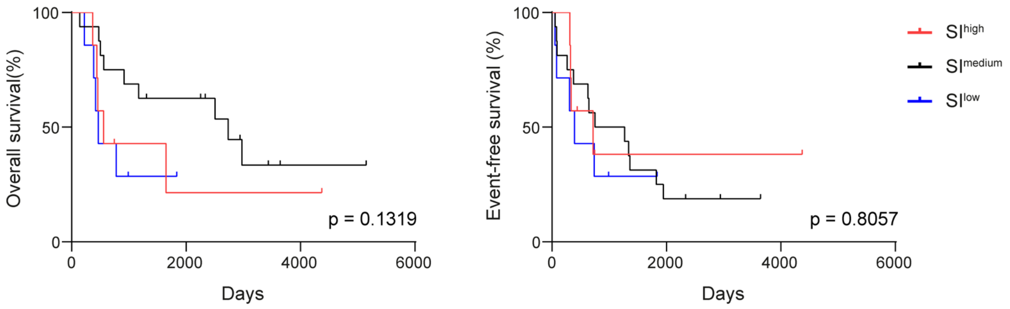


**Figure S4: Kaplan-Meier analysis of sarcoma patients stratified by SI values of the respective tumor-derived cell lines.** SI stratification was performed relative to median: $SI^{high}$, upper quartile; $SI^{lmedium}$, interquartile range; $SI^{low}$, lower quartile. Statistical significance was determined by Mantel-Cox test.

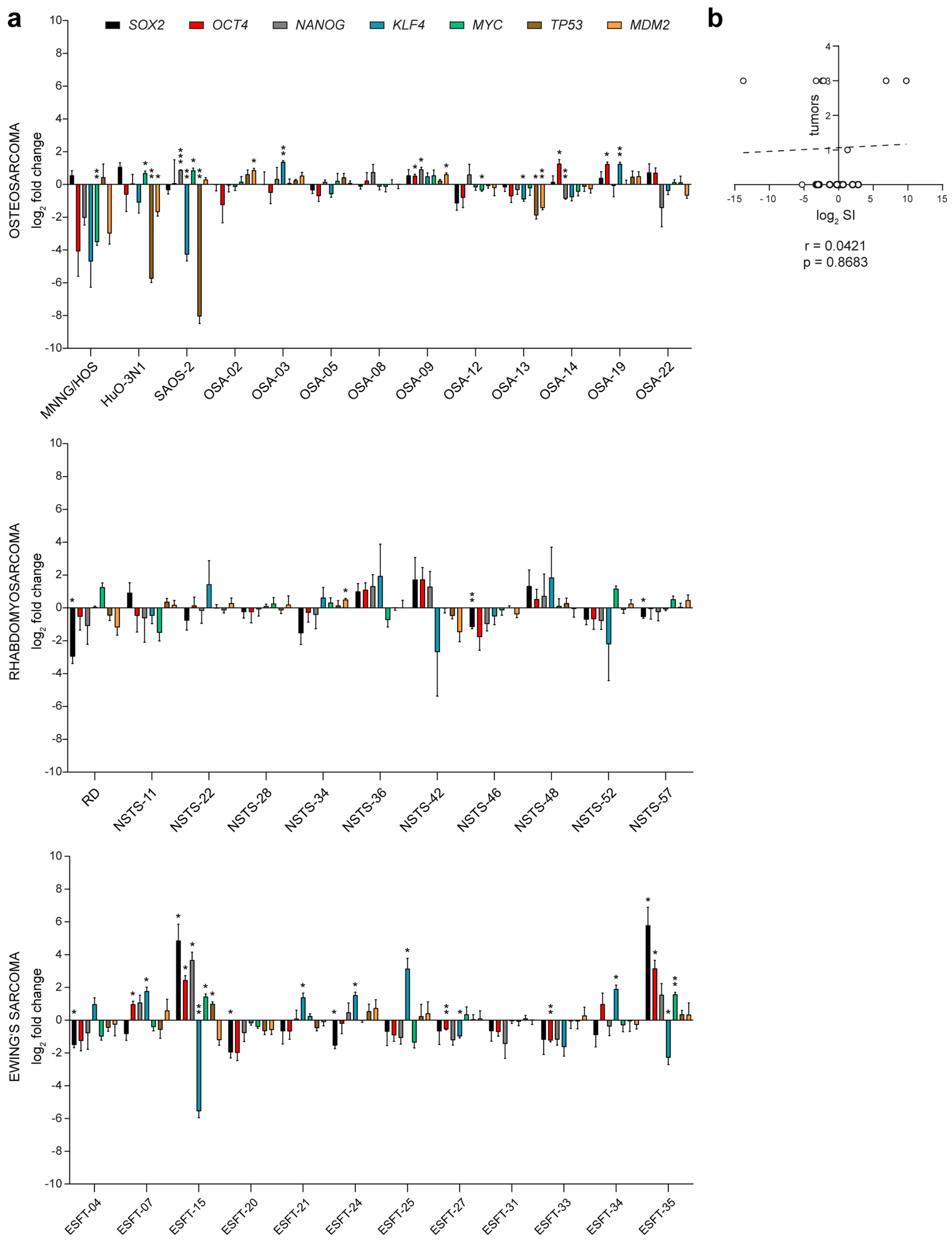


**Figure S5: qPCR analysis of mRNA expression of selected stemness-associated TFs, *TP53* and *MDM2* in a panel of OS, RMS and ES established and patient-derived cell lines. a)** Expression values of each cell line were evaluated relative to the average mRNA expression value of the respective gene in each subgroup of patient-derived sarcomas. Data are mean ± SD, biological n=3. **b)** Correlation analysis between the SI value calculated from qPCR data ($\log_2$) and the number of xenografts derived from selected $SI^{high}$ and $SI^{low}$ OS, RMS and ES cell lines. Statistical significance was determined by one-way ANOVA with Welch's correction followed by post-hoc Dunnett's test (**a**), r was determined by Spearman correlation test (**b**), *p<0.05, **p<0.01, ***p<0.001.

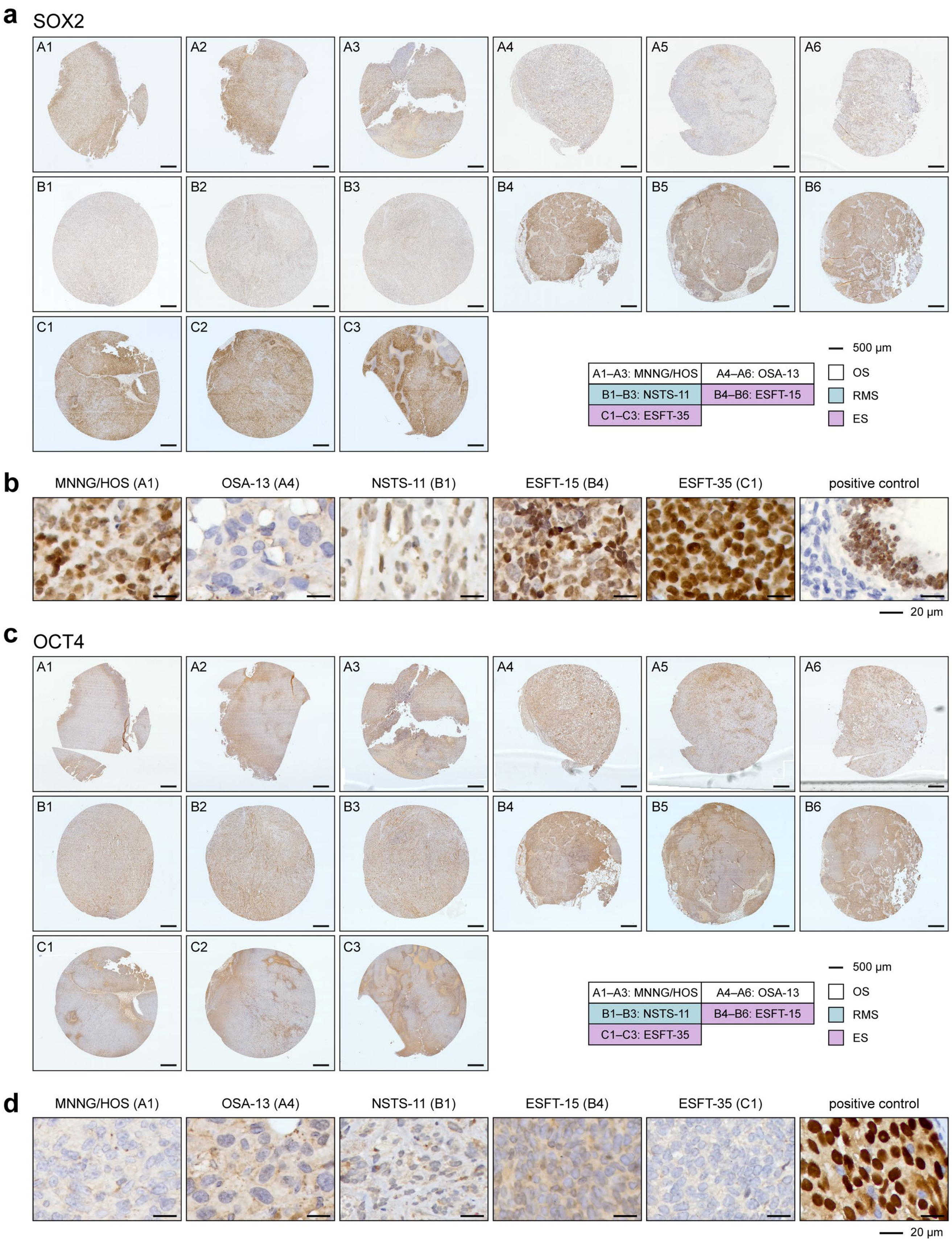


**Figure S6: IHC analysis of SOX2 and OCT4 in sarcoma xenograft tissues. a,c)** SOX2 (a) and OCT4 (c) expression in the whole scanned microarray cores of xenograft tissues. **b,d)** Representative close-up images of SOX2 (b) and OCT4 (d) IHC detection in selected xenografts and a positive control tissues, fetal lungs and seminoma, respectively.

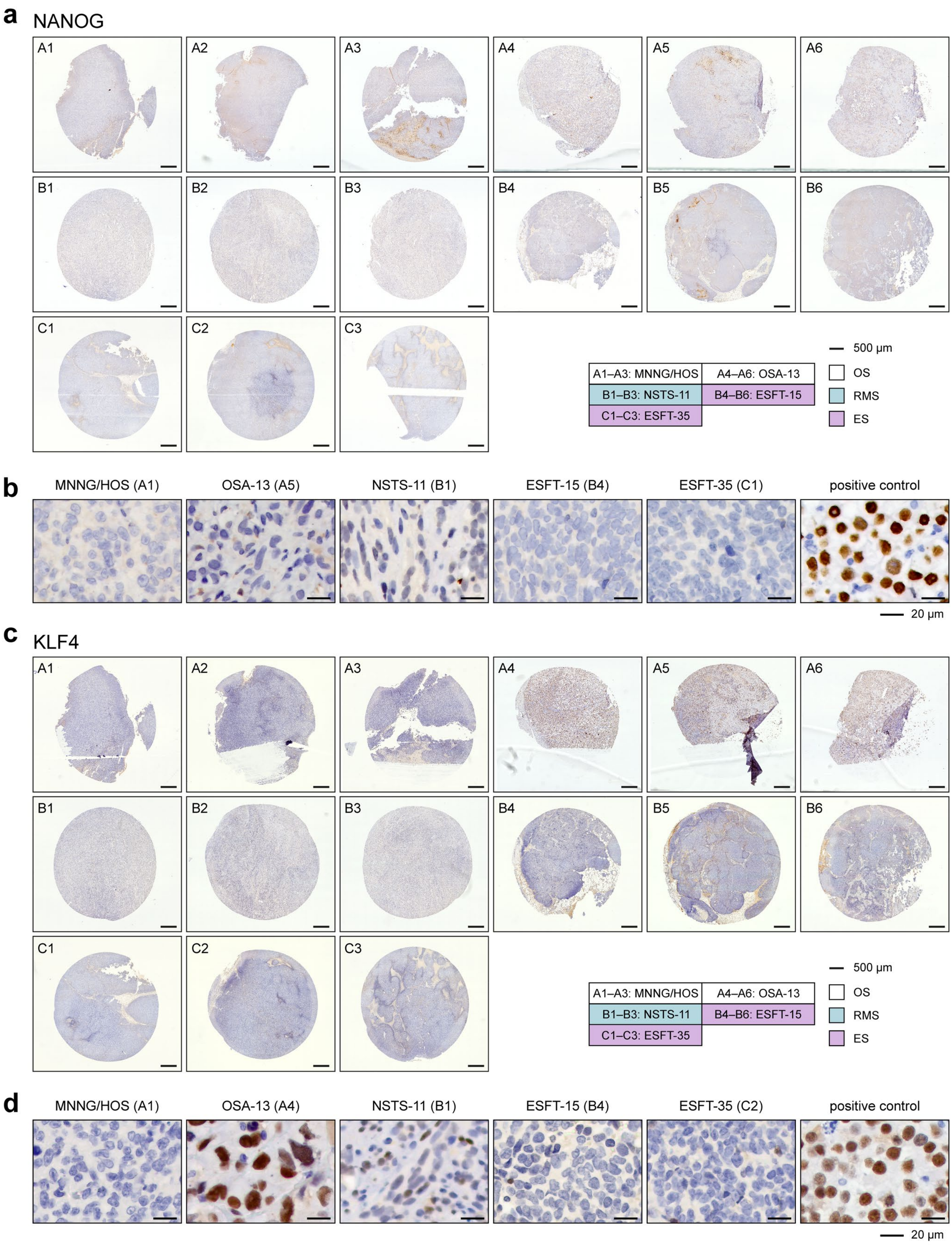


**Figure S7: IHC analysis of NANOG and KLF4 in sarcoma xenograft tissues. a,c)** NANOG (a) and KLF4 (c) expression in the whole scanned microarray cores of xenograft tissues. **b,d)** Representative close-up images of NANOG (b) and KLF4 (d) IHC detection in selected xenografts and a positive control seminoma tissue.

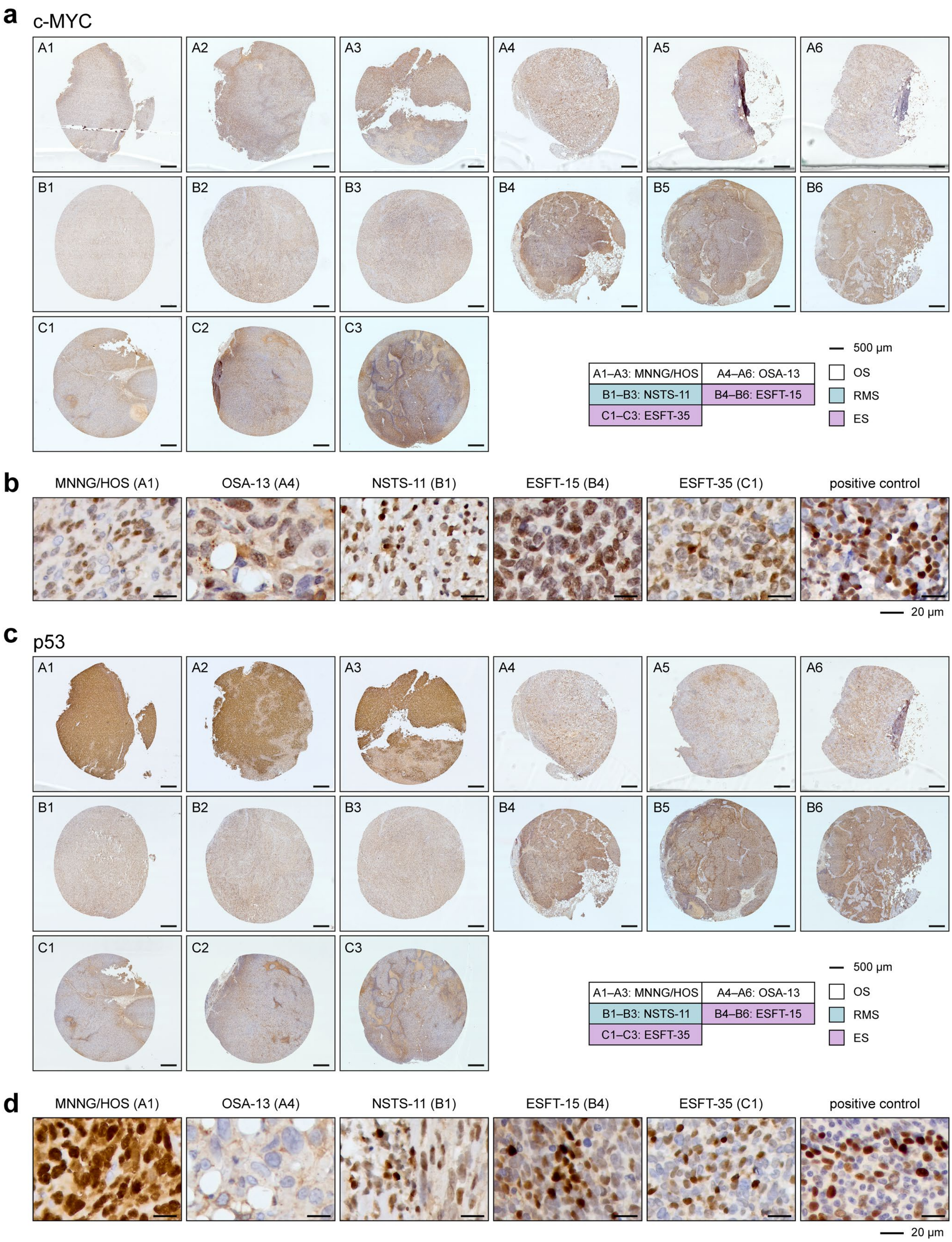


**Figure S8: IHC analysis of c-MYC and p53 in sarcoma xenograft tissues. a,c)** c-MYC (a) and p53 (c) expression in the whole scanned microarray cores of xenograft tissues. **b,d)** Representative close-up images of c-MYC (b) and p53 (d) IHC detection in selected xenografts and a positive control tissues, Burkitt lymphoma and tonsilla, respectively.

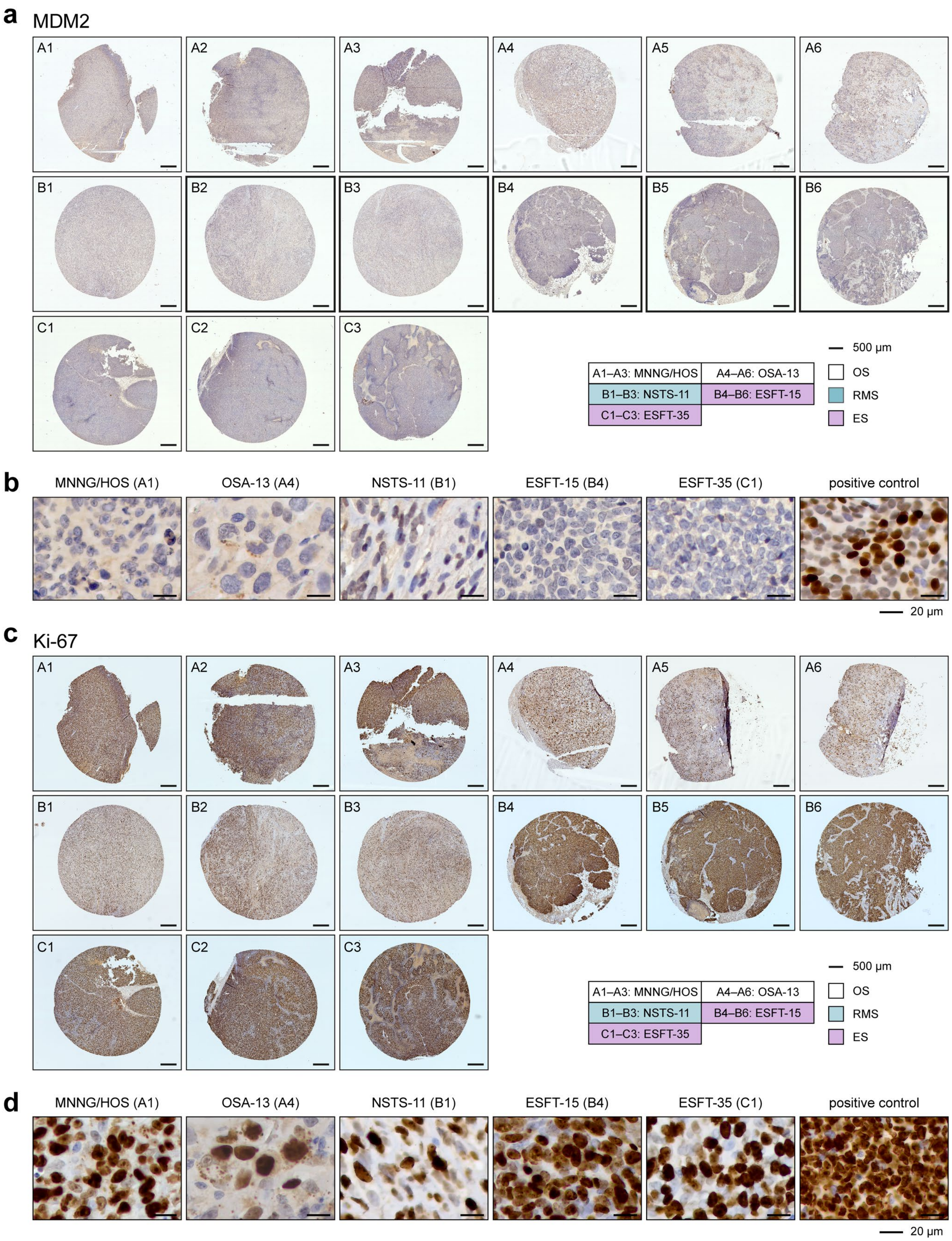


**Figure S9: IHC analysis of MDM2 and Ki-67 in sarcoma xenograft tissues. a,c)** MDM2 (a) and Ki-67 (c) expression in the whole scanned microarray cores of xenograft tissues. **b,d)** Representative close-up images of MDM2 (b) and Ki-67 (d) IHC detection in selected xenografts and a positive control tissues, undifferentiated epithelioid sarcoma and tonsilla, respectively.

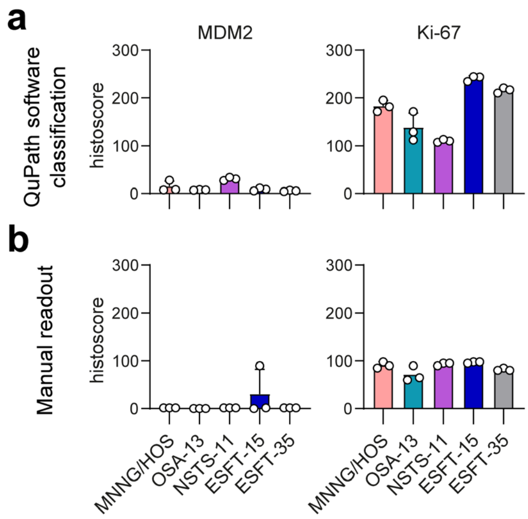


**Figure S10: Histoscore analysis of MDM2 and Ki-67 in sarcoma xenografts. a,b)** IHC staining of MDM2 and Ki-67 assessed computationally using QuPath software (a) and by manual readout by experienced pathologist (b), biological n=3.

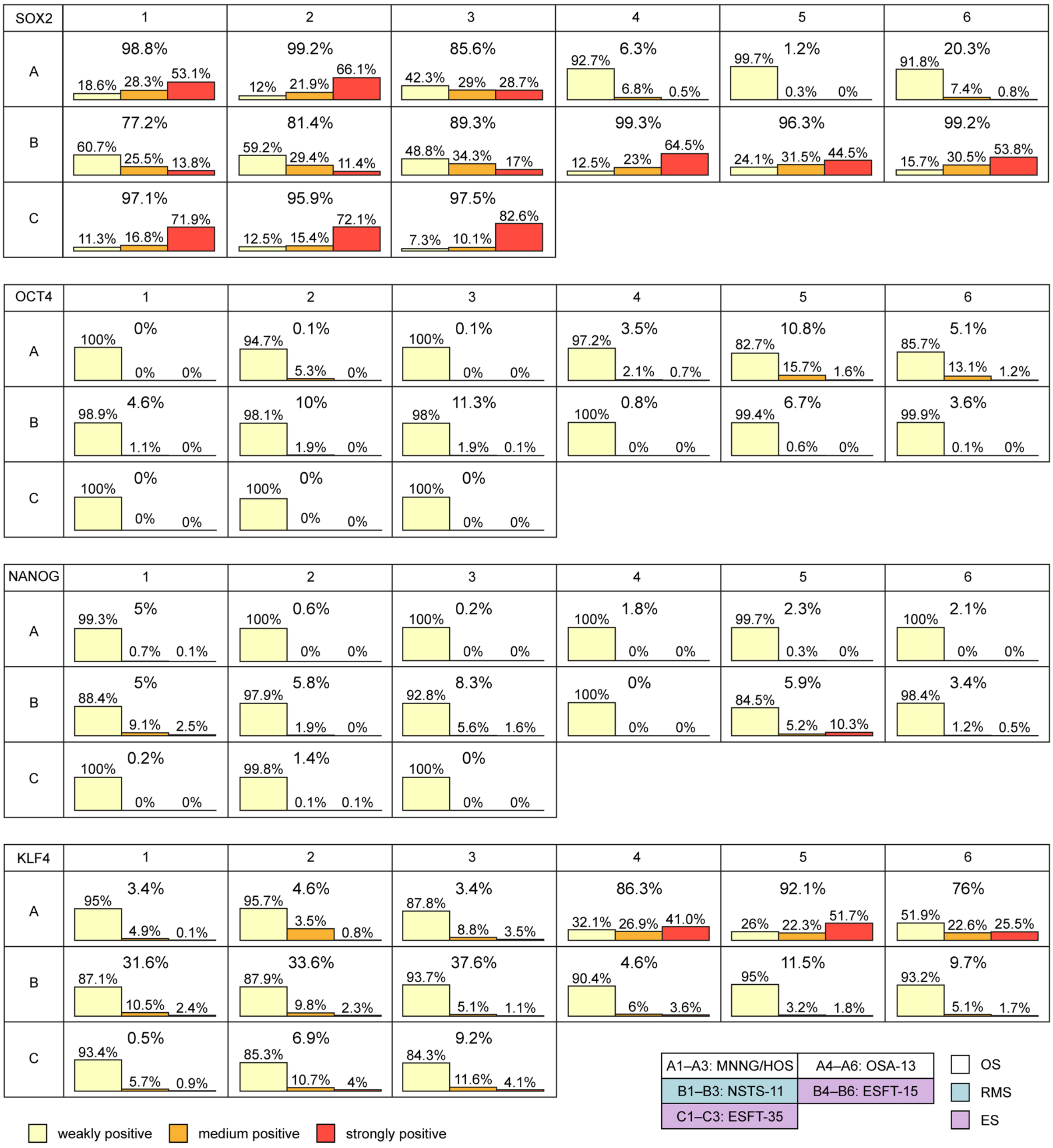


**Figure S11: Detailed computational evaluation of SOX2, OCT4, NANOG and KLF4 staining in sarcoma xenografts.** Immunoreactivity and percentage of positive cells as determined by QuPath software following the IHC staining.

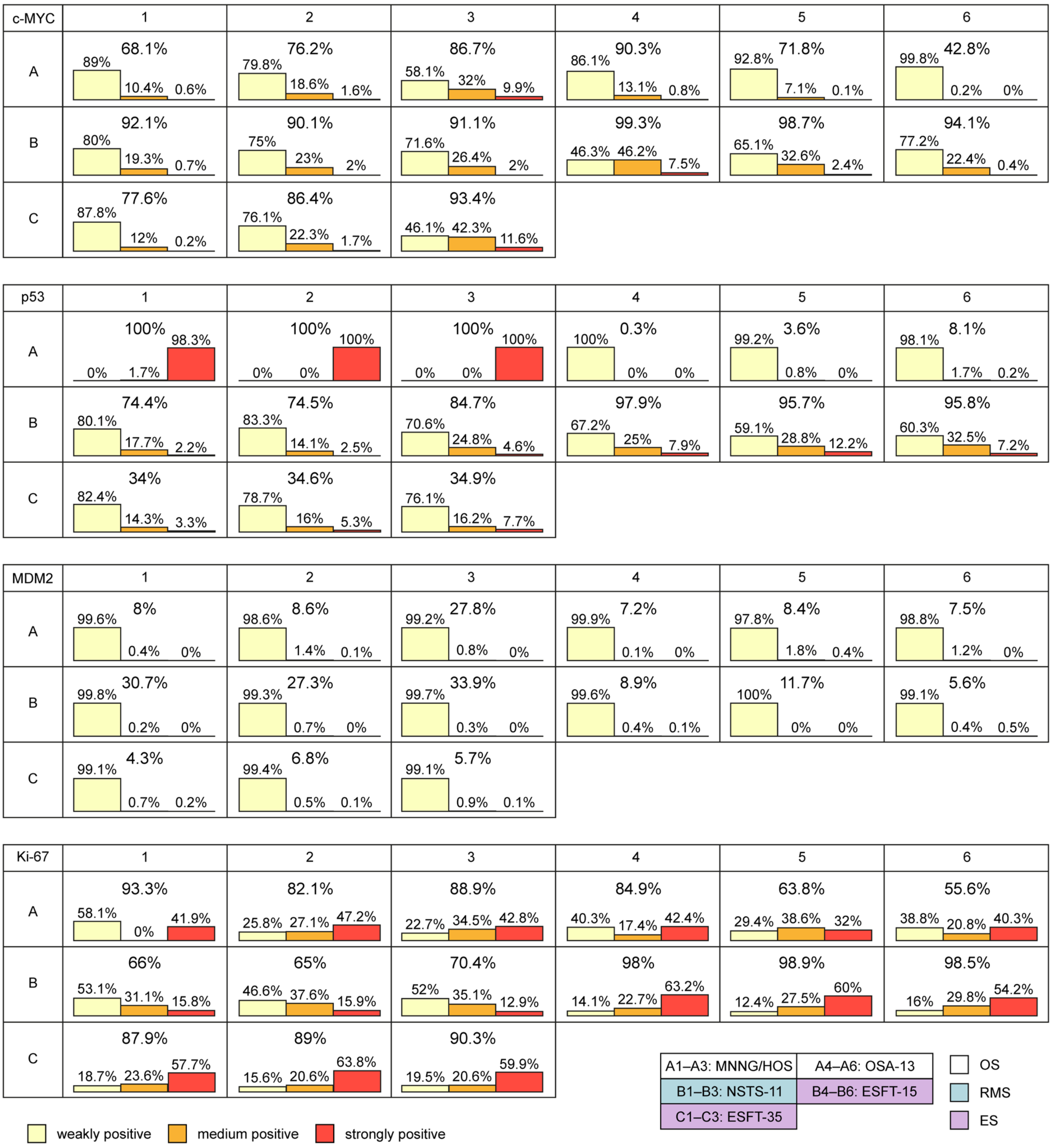


**Figure S12: Detailed computational evaluation of c-MYC, p53, MDM2 and Ki-67 staining in sarcoma xenografts.** Immunoreactivity and percentage of positive cells as determined by QuPath software following the IHC staining.

| SOX2 | 1 | 2 | 3 | 4 | 5 | 6 |
|---|---|---|---|---|---|---|
| A | 100% (+++) | 100% (+++) | 60% (++) | 0% | 0% | 0% |
| B | 100% (+/++) | 100% (+/++) | 90% (++) | 100% (++/+++) | 100% (++/+++) | 100% (++) |
| C | 100% (+++) | 100% (+++) | 100% (+++) | | | |

| OCT4 | 1 | 2 | 3 | 4 | 5 | 6 |
|---|---|---|---|---|---|---|
| A | 0% | 0% | 0% | 0% | 0% | 0% |
| B | 10% (++) | 20% (++) | 15% (++) | <5% (++) | <5% (++) | <5% (++) |
| C | 0% | 0% | <5% (++) | | | |

| NANOG | 1 | 2 | 3 | 4 | 5 | 6 |
|---|---|---|---|---|---|---|
| A | 0% | 0% | 0% | 0% | 5% (+++) | 0% |
| B | <1% (+++) | <1% (+++) | <1% (+++) | 0% | <1% (+++) | <1% (+++) |
| C | 0% | 0% | 0% | | | |

| KLF4 | 1 | 2 | 3 | 4 | 5 | 6 |
|---|---|---|---|---|---|---|
| A | 0% | <1% (+++) | 0% | 80% (+++) | 90% (+++) | 80% (+++) |
| B | 5% (+/++) | 10% (+/++) | 5% (+/++) | 0% | <1% (++) | 0% |
| C | <1% (+++) | <1% (++) | <1% (++) | | | |

| c-MYC | 1 | 2 | 3 | 4 | 5 | 6 |
|---|---|---|---|---|---|---|
| A | 70% (++) | 80% (++/+++) | 80% (++/+++) | 30% (++) | 40% (++) | <1% (++) |
| B | 100% (+++) | 90% (++/+++) | 80% (++/+++) | 90% (+++) | 90% (+++) | 90% (+++) |
| C | 70% (++) | 60% (++) | 70% (+++) | | | |

| p53 | 1 | 2 | 3 | 4 | 5 | 6 |
|---|---|---|---|---|---|---|
| A | 100% | 100% | 100% | 0% | 0% | 0% |
| B | 85% | 85% | 90% | 90% | 80% | 90% |
| C | 50% | 55% | 50% | | | |

| MDM2 | 1 | 2 | 3 | 4 | 5 | 6 |
|---|---|---|---|---|---|---|
| A | <1% (++) | <1% (++) | <1% (++) | 0% | 0% | 0% |
| B | 1% (++) | 1% (++) | 1% (++) | 0% | <1% (++) | 90% |
| C | <1% (++) | <1% (++) | 1% (++) | | | |

| Ki-67 | 1 | 2 | 3 | 4 | 5 | 6 |
|---|---|---|---|---|---|---|
| A | 98% | 90% | 85% | 90% | 60% | 65% |
| B | 95% | 90% | 95% | 98% | 98% | 95% |
| C | 85% | 80% | 80% | | | |

| | | | |
|---|---|---|---|
| A1–A3: MNNG/HOS | A4–A6: OSA-13 | | OS |
| B1–B3: NSTS-11 | B4–B6: ESFT-15 | | RMS |
| C1–C3: ESFT-35 | | | ES |

**Figure S13: Manual readout of the IHC staining of selected stemness-associated and p53 pathway-related proteins in sarcoma xenografts.** Immunoreactivity and percentage of positive cells determined by experienced pathologist following the IHC staining of the respective proteins in xenograft tissues formed by the indicated sarcoma cell lines.

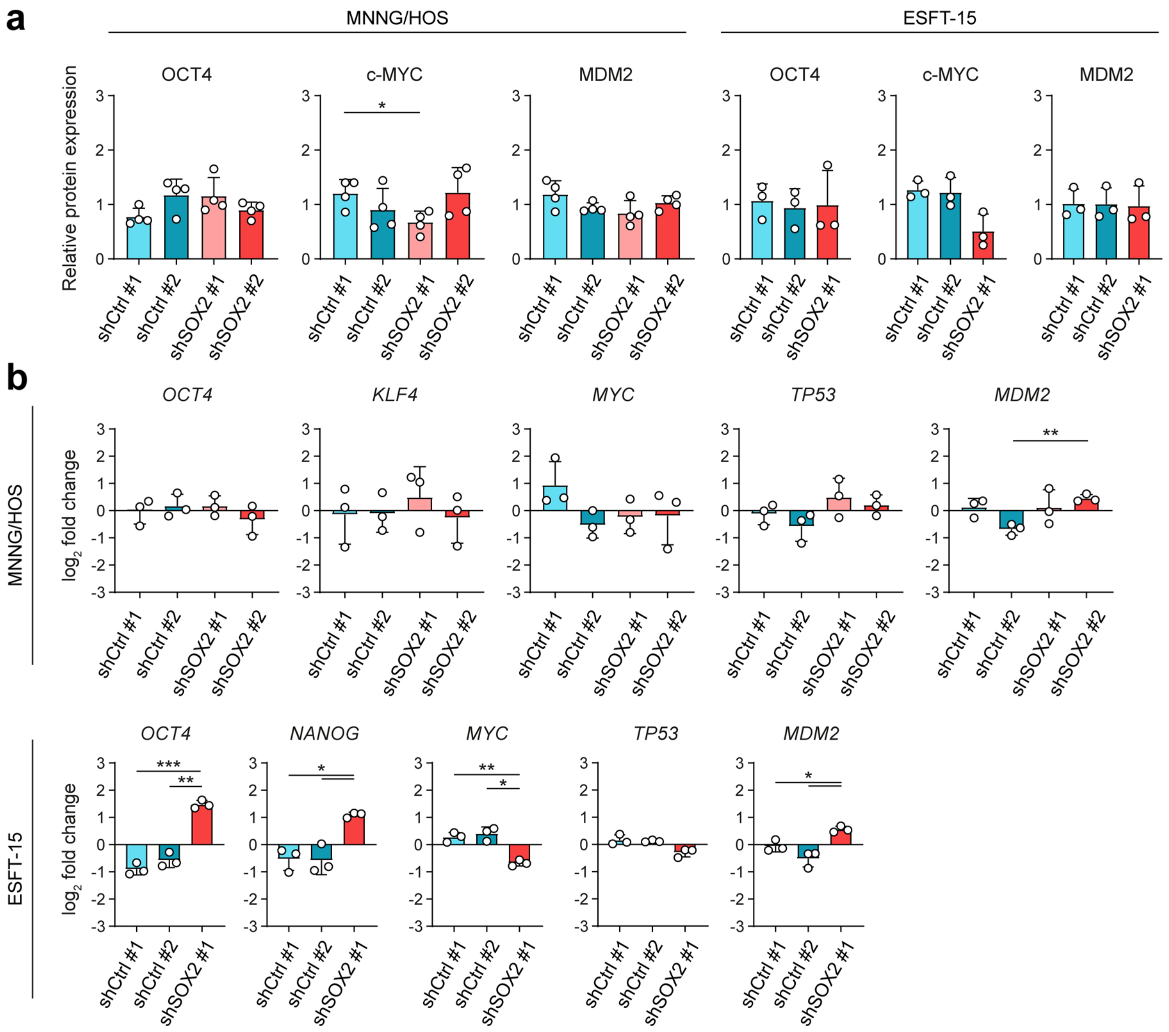


**Figure S14: Expression analysis of stemness-associated TFs, p53 and MDM2 in MNNG/HOS- and ESFT-15-derived shCtrl and shSOX2 single-cell clones. a)** Densitometric analysis of the protein levels. **b)** mRNA expression analyzed by qPCR. Data presented as mean ± SD, biological n=3–4. Statistical significance was determined by one-way ANOVA with Welch's correction followed by post-hoc Dunnett's test, *p<0.05, **p<0.01, ***p<0.001.

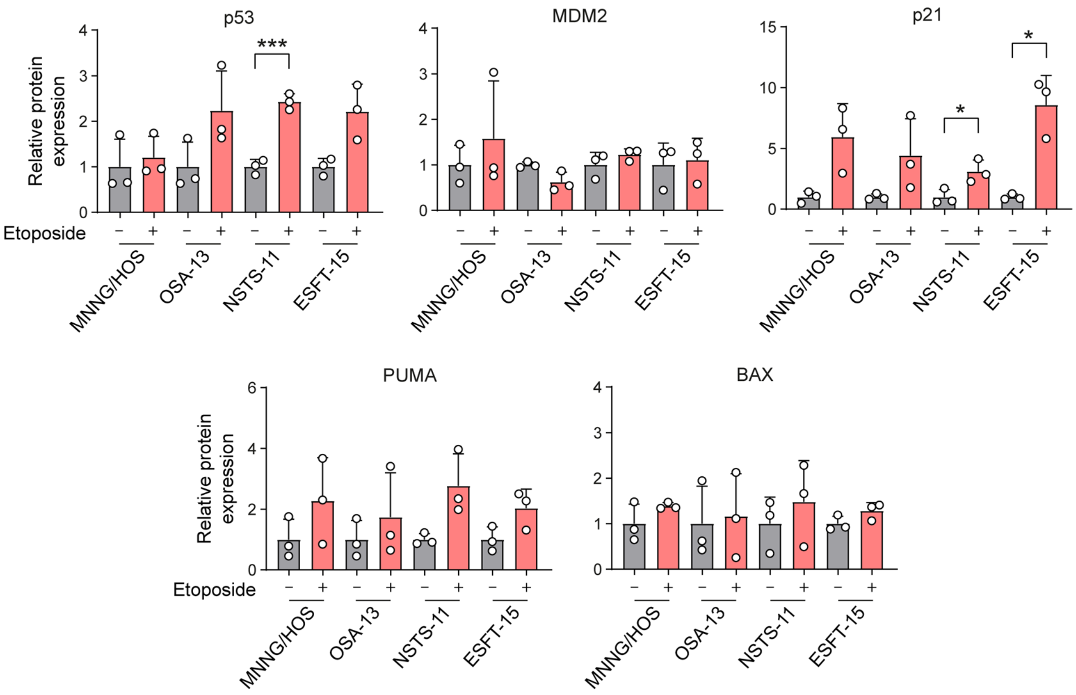


**Figure S15: Densitometric analysis of p53 and its downstream targets in mut-p53 MNNG/HOS and wt-p53 SI$^{high}$ cell lines after 24-h treatment by 25µM etoposide.** Statistical significance was determined by Welch's t-test, biological n=3, *p<0.05, ***p<0.001.

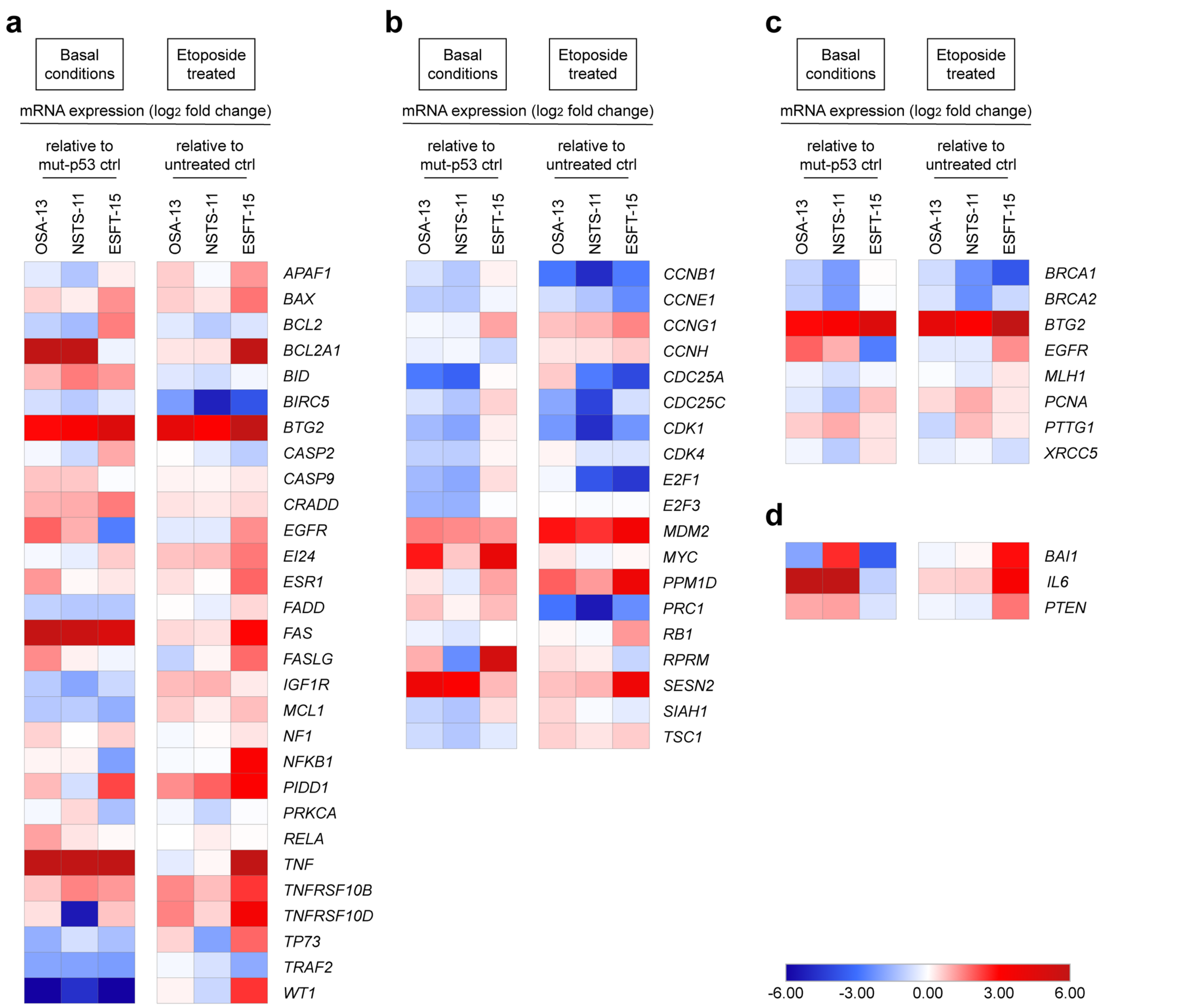


**Figure S16: qPCR analysis of mRNA expression of downstream genes regulated by p53 in selected SI$^{high}$ sarcoma cell lines.** Analyzed genes are involved in **(a)** apoptosis, **(b)** cell cycle regulation, **(c)** DNA damage and repair and **(d)** angiogenesis**.** mRNA expression of each gene was analyzed at basal level compared with the expression of respective gene in MNNG/HOS cell line (left panel) and after 24-h treatment by 25µM etoposide (right panel), biological n=1.

**Supplementary Table 1: Patient-derived osteosarcoma, rhabdomyosarcoma and Ewing's sarcoma cell lines and clinicopathological characteristics of the respective cases**

| Sarcoma subtype | Cell line | Gender | Age | Site | Post therapy | Fusion gene[a] |
|---|---|---|---|---|---|---|
| OS | OSA-02 | M | 20 | primary | no | - |
| | OSA-03 | M | 14 | primary | no | - |
| | OSA-05 | M | 9 | primary | no | - |
| | OSA-08 | M | 10 | primary | no | - |
| | OSA-09 | F | 22 | primary | no | - |
| | OSA-12 | M | 17 | primary | no | - |
| | OSA-13 | F | 6 | primary | no | - |
| | OSA-14 | F | 16 | primary | no | - |
| | OSA-19 | M | 2 | primary | no | - |
| | OSA-22 | M | 9 | primary | no | - |
| RMS | NSTS-11 | F | 16 | primary | yes | n/a |
| | NSTS-22 | F | 6 | primary | no | *PAX3/FOXO1* |
| | NSTS-28 | M | 8 | primary | no | negative |
| | NSTS-34 | M | 3 | primary | no | negative |
| | NSTS-36 | F | 2 | primary | yes | *PAX3/FOXO1* |
| | NSTS-42 | F | 13 | primary | no | *PAX3/FOXO1* |
| | NSTS-46 | M | 5 | primary | no | negative |
| | NSTS-48 | F | 7 | primary | yes | *PAX3/FOXO1* |
| | NSTS-52 | M | 4 | primary | yes | negative |
| | NSTS-57 | M | 15 | metastasis | yes | *PAX3/FOXO1* |
| ES | ESFT-04 | M | 3 | primary | no | *EWSR1/FLI1* |
| | ESFT-07 | M | 15 | primary | no | *EWSR1/ERG* |
| | ESFT-15 | M | 17 | primary | no | *EWSR1/FLI1* |
| | ESFT-20 | M | 10 | primary | no | *EWSR1/FLI1* |
| | ESFT-21 | M | 18 | primary | no | *EWSR1/FLI1* |
| | ESFT-24 | M | 16 | primary | no | *EWSR1/FLI1* |
| | ESFT-25 | F | 13 | metastasis | yes | *EWSR1/FLI1* |
| | ESFT-27 | F | 24 | primary | yes | *EWSR1* break |
| | ESFT-31 | M | 12 | primary | no | *EWSR1/FLI1* |
| | ESFT-33 | M | 12 | primary | no | *EWSR1/FLI1* |
| | ESFT-34 | F | 14 | primary | no | *EWSR1* break |
| | ESFT-35 | F | 20 | metastasis | yes | *EWSR1/FLI1* |

[a] Sarcoma subtype-specific translocations detected in the respective tumor tissue; n/a, not available; negative, no *PAX3/7* translocation. OS, osteosarcoma; RMS, rhabdomyosarcoma; ES, Ewing's sarcoma.

**Supplementary Table 2: Cell lines and their culture media components**

| Cell type | Cell line | Culture medium | Supplements |
|---|---|---|---|
| OS, RMS, ES | patient-derived cell lines | DMEM Low Glucose (LM-D1100, Biosera) | 20% FBS (FB-1101); 2 mM L-Glutamine (XC-T1715); Penicilin (100 IU/ml) /Streptomycin (100 μg/ml) (X-A4122) all Biosera |
| OS | HuO-3N1 | | |
| | SAOS-2 | | 10% FBS (FB-1101); 2 mM L-Glutamine (XC-T1715); Penicilin (100 IU/ml) /Streptomycin (100 μg/ml) (X-A4122) all Biosera |
| | MNNG/HOS | | 10% FBS (FB-1101); 2 mM L-Glutamine (XC-T1715); 1× MEM Non-Essential Amino Acids (XC-E1154); Penicilin (100 IU/ml) /Streptomycin (100 μg/ml) (XC-A4122) all Biosera |
| embryonal carcinoma | NTERA-2 | DMEM High Glucose (LM-D1112, Biosera) | 10% FBS (FB-1101); 2 mM L-Glutamine (XC-T1715); Penicilin (100 IU/ml) /Streptomycin (100 μg/ml) (X-A4122) all Biosera |
| RMS | RD | | 10% FBS (FB-1101); 2 mM L-Glutamine (XC-T1715); 1× MEM Non-Essential Amino Acids (XC-E1154); Penicilin (100 IU/ml) /Streptomycin (100 μg/ml) (X-A4122) all Biosera |
| human neonatal dermal fibroblasts | NDF-2 and NDF-3 | | |

Provider: Biosera (Nuaille, France). FBS, fetal bovine serum; OS, osteosarcoma; RMS, rhabdomyosarcoma; ES, Ewing’s sarcoma.

**Supplementary Table 3: Primer sequencies used for qPCR**

| | | |
|---|---|---|
| *SOX2* | forward (5’→3’) | ACATGAACGGCTGGAGCAA |
| | reverse (5’→3’) | GTAGGACATGCTGTAGGTGGG |
| *POU5F1* | forward (5’→3’) | TGGAGAAGGAGAAGCTGGAGCAAAA |
| | reverse (5’→3’) | GGCAGATGGTCGTTTGGCTGAATA |
| *NANOG* | forward (5’→3’) | AATACCTCAGCCTCCAGCAGAT |
| | reverse (5’→3’) | TGCGTCACACCATTGCTATTCTTC |
| *KLF4* | forward (5’→3’) | ATCTTTCTCCACGTTCGCGTCTG |
| | reverse (5’→3’) | AAGCACTGGGGGAAGTCGCTTC |
| *MYC* | forward (5’→3’) | TCTCTCCGTCCTCGGATTCT |
| | reverse (5’→3’) | GCCTCTTTTCCACAGAAACAACA |
| *TP53* | forward (5’→3’) | GCACTGGTGTTTTGTTGTGG |
| | reverse (5’→3’) | GTGGTTTCAAGGCCAGATGT |
| *MDM2* | forward (5’→3’) | GGCCTGCTTTACATGTGCAA |
| | reverse (5’→3’) | GCACAATCATTTGAATTGGTTGTC |
| *HSP90AB1* | forward (5’→3’) | CGCATGAAGGAGACACAGAA |
| | reverse (5’→3’) | TCCCATCAAATTCCTTGAGC |

**Supplementary Table 4: Antibodies and detection reagents for immunohistochemistry**

| Antibody, host, type [clone] | Cell conditioning | Dilution | Primary antibody incubation | Detection | Positive control |
|---|---|---|---|---|---|
| SOX2, Rb, mAb [EPR3131] | 100 °C, 72 min | 1:100 | 44 min | Opti | fetal lungs |
| OCT4, Rb pAb | 95 °C, 48 min | 1:500 | 32 min | Opti | seminoma |
| NANOG, Rb mAb [EPR2027(2)] | 95 °C, 52 min | 1:200 | 32 min | Ultra | seminoma |
| KLF4, Rb, mAb [EPR19590] | 95 °C, 52 min | 1:2000 | 32 min | Ultra | seminoma |
| c-MYC, Rb, mAb [Y69] | 98 °C, 40 min | 1:50 | 24 min | Opti | Burkitt lymphoma |
| p53, Mo, mAb [DO-7] | 100 °C, 72 min | - | 28 min | Opti | tonsilla |
| MDM2, Mo, mAb [3G187] | 95 °C, 52 min | 1:50 | 40 min | Ultra | undifferentiated epithelioid sarcoma |
| Ki-67, Rb, mAb [SP6] | 100 °C, 72 min | 1:200 | 28 min | Opti | tonsilla |

Rb, rabbit; Mo, mouse; mAb, monoclonal antibody; pAb, polyclonal antibody; Opti, OptiView DAB IHC Detection Kit (Roche); Ultra, UltraView Universal DAB Detection Kit (Roche).
